# Early Visual Cortex Represents Sensory and Mnemonic Orientations in Separate Subspaces with Preserved Geometry

**DOI:** 10.64898/2026.04.20.718367

**Authors:** Sungje Kim, Jaeseob Lim, Hyunwoo Gu, Heeseung Lee, Hyang-Jung Lee, Minjin Choe, Dong-gyu Yoo, Joonwon Lee, Sang-Hun Lee

## Abstract

How early visual cortex (EVC) represents working memory (WM) content while continuing to process incoming sensory input remains unclear. Using fMRI with prolonged delays to isolate mnemonic activity, together with analyses of cross-decoding, low-dimensional subspace structure, and representational geometry, we examined the relationship between sensory and mnemonic orientation representations in human EVC. Cross-decoding generalized poorly between sensory and mnemonic epochs, but this did not imply unrelated codes. Rather, the two occupied separable low-dimensional subspaces while preserving representational geometry across epochs. During discrimination and estimation, sensory- and mnemonic-trained decoders yielded dissociable readouts of concurrent sensory and mnemonic information from the same EVC measurements. Mnemonic coding showed little dependence on the retinotopic radial bias that characterized sensory coding, and trial-by-trial variability in mnemonic representation predicted both discrimination choices and estimation reports. Our findings support a population-level account in which mnemonic information in EVC is re-expressed in a separable but geometrically preserved format.

## Introduction

Constantly immersed in fleeting sensory inputs, the brain must flexibly combine or differentiate past and current information to support stable perception and adaptive behavior. When a deer dashes through a dense forest, we piece together a unified percept from fragmented glimpses. In a grocery aisle, we compare items no longer in view with those currently before us. At the core of these abilities is working memory (WM): the capacity to maintain and use information—some from the past, others newly arriving—within what we experience as an “extended present” (**James, 1890**^1^). How does the brain support this feat?

For vision, one prominent answer has been the ‘sensory recruitment’ hypothesis: feature-selective activity patterns in early visual cortex (EVC) during sensory encoding are re-engaged during memory delays (**D’Esposito & Postle, 2015**^2^). Indeed, numerous human neuroimaging (**Harrison & Tong, 2009**^3^**; Serences et al., 2009**^4^**; Riggall & Postle, 2012**^5^**; Christophel et al., 2017**^6^**; Curtis & Sprague, 2021**^7^) and monkey electrophysiology (**Huang et al., 2024**^8^) studies reveal that stimulus features can be reliably decoded from EVC during blank delays. However, this raises a fundamental paradox: how does EVC concurrently process new sensory inputs without catastrophic interference with maintained memory representations (**Mendoza-Halliday et al., 2014**^9^**; Stokes, 2015**^10^**; Bettencourt & Xu, 2016**^11^**; Xu, 2017**^12^**, 2020**^13^), and how are these two information streams differentiated yet related within the same cortical population? The central question is therefore no longer simply whether mnemonic information is present in EVC, but whether sensory recruitment can provide a viable account of WM when remembered and incoming sensory information must coexist within the same cortical population (**Kiyonaga & Serences, 2025; Grassi et al., 2024; Phylactou et al., 2024**).

To address this issue, we must characterize the relationship between sensory and mnemonic representations in population activity space (**Jazayeri & Afraz, 2017**^14^**; Kriegeskorte & Wei, 2021**^15^**; Saxena & Cunningham, 2019**^16^). At this level, three broad scenarios are possible (**Fig. 1a-c**). First, sensory and mnemonic signals could share a common population code, occupying the same subspace of population activity and exhibiting similar representational geometry (**Fig. 1a**). Second, they could occupy separate subspaces and differ in representational geometry, consistent with distinct sensory and mnemonic population codes (**Fig. 1b**). Third, they could occupy separate subspaces while preserving their internal geometry across conditions—a relationship we term the re-embedded-code scenario (**Fig. 1c**). In this case, sensory and mnemonic information would preserve their representational geometry while being expressed along different axes of the measured population activity.

**Figure 1.**
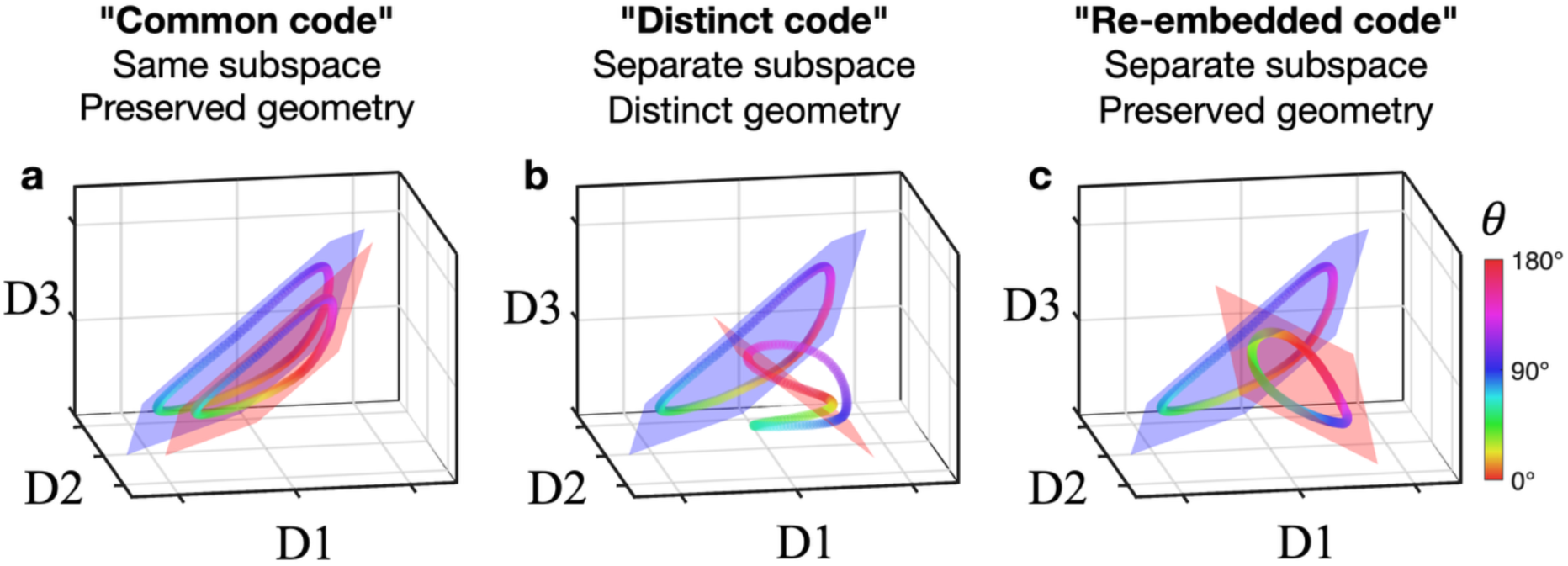
| Conceptual scenarios for sensory and mnemonic population codes in EVC. a–c, Three theoretical scenarios illustrating relationships between sensory and mnemonic population codes. Schematic population-activity manifolds are shown in a three-dimensional population state space (axes: three response dimensions), color-coded by orientation *θ* (see legend). Translucent planes indicate the subspace associated with each manifold (sensory, blue; mnemonic, red). **a,** *Common code*. Sensory and mnemonic signals occupy the same population subspace and share the same representational geometry. **b,** *Distinct code*. Sensory and mnemonic signals occupy separate subspaces and differ in representational geometry. **c,** *Re-embedded code*. Sensory and mnemonic signals occupy separate subspaces while preserving a shared representational geometry.

Distinguishing among these scenarios is important because they have different implications for two computational demands posed by the paradox: limiting interference between past and present information, and enabling their direct comparison or combination when behavior requires it. Subspace separation should, in principle, reduce mutual interference by allowing sensory and mnemonic signals to coexist in separable population dimensions. Geometry preservation, by contrast, should facilitate comparison or combination by maintaining a compatible representational format across states. The common-code scenario naturally supports format compatibility but offers less protection from interference; the distinct-code scenario favors resistance to interference but not direct compatibility; only the re-embedded-code scenario potentially satisfies both demands at once. Determining which of these representational relationships best characterizes EVC is therefore central to evaluating sensory-recruitment accounts of WM.

Prior work has not clearly distinguished among these scenarios, in part because the dominant analytical tools—especially decoding and cross-decoding applied to temporally overlapping epochs—do not uniquely adjudicate their representational relationships (**Sandhaeger & Siegel, 2023**). In particular, a failure of cross-decoding does not by itself imply a distinct-code scenario: it can arise either because sensory and mnemonic signals truly differ in representational geometry or because a shared geometry has been re-expressed in a different population subspace through drift or rotation in population space (**King & Dehaene, 2014**^17^**; Naselaris & Kay, 2015**^18^**; Murray et al., 2017**^19^). Conversely, successful cross-decoding in fMRI studies with short delays can be difficult to interpret because lingering stimulus-evoked activity may contaminate nominal delay-period signals, making active mnemonic maintenance hard to distinguish from a passive fading echo of sensory input (**Supplementary Fig. 1**). In EVC, this ambiguity is compounded by coarse-scale spatial biases in fMRI measurements, such as retinotopic radial bias, which can create spurious similarities in decoding structure across epochs (**Sandhaeger & Siegel, 2023**^20^). As a result, prior work has left unresolved the representational relationship between sensory and mnemonic codes in EVC.

To address these limitations, we used fMRI to temporally isolate late mnemonic activity from stimulus-locked responses (**Fig. 2a**) and combined this design with population-level analyses of dimensionality reduction, representational geometry, and decoding (**Kaufman et al., 2014**^21^**; Cunningham & Yu, 2014**^22^**; Elsayed et al., 2016**^23^**; Murray et al., 2017**^19^**; Kobak et al., 2016**^24^**; Kriegeskorte & Wei,2021**^15^**; Jazayeri & Ostojic, 2021**^25^). We found that poor cross-decoding between sensory and mnemonic epochs did not imply unrelated codes. Rather, sensory and mnemonic orientation representations in human EVC occupied separable low-dimensional subspaces while preserving ring-like orientation structure across epochs. Consistent with this re-embedded relationship, sensory- and mnemonic-trained decoders yielded dissociable readouts of concurrently relevant sensory and mnemonic information during discrimination and estimation. We further found that, whereas sensory representations expressed retinotopic radial bias, mnemonic coding showed little dependence on that bias, indicating a decoupling of mnemonic orientation coding from the coarse retinotopic bias that characterizes sensory encoding. Finally, trial-by-trial variability in the mnemonic representation predicts both discrimination choices and estimation reports. Together, these findings support a population-level account in which mnemonic information in EVC is re-expressed in a separable but geometrically preserved format.

**Figure 2.**
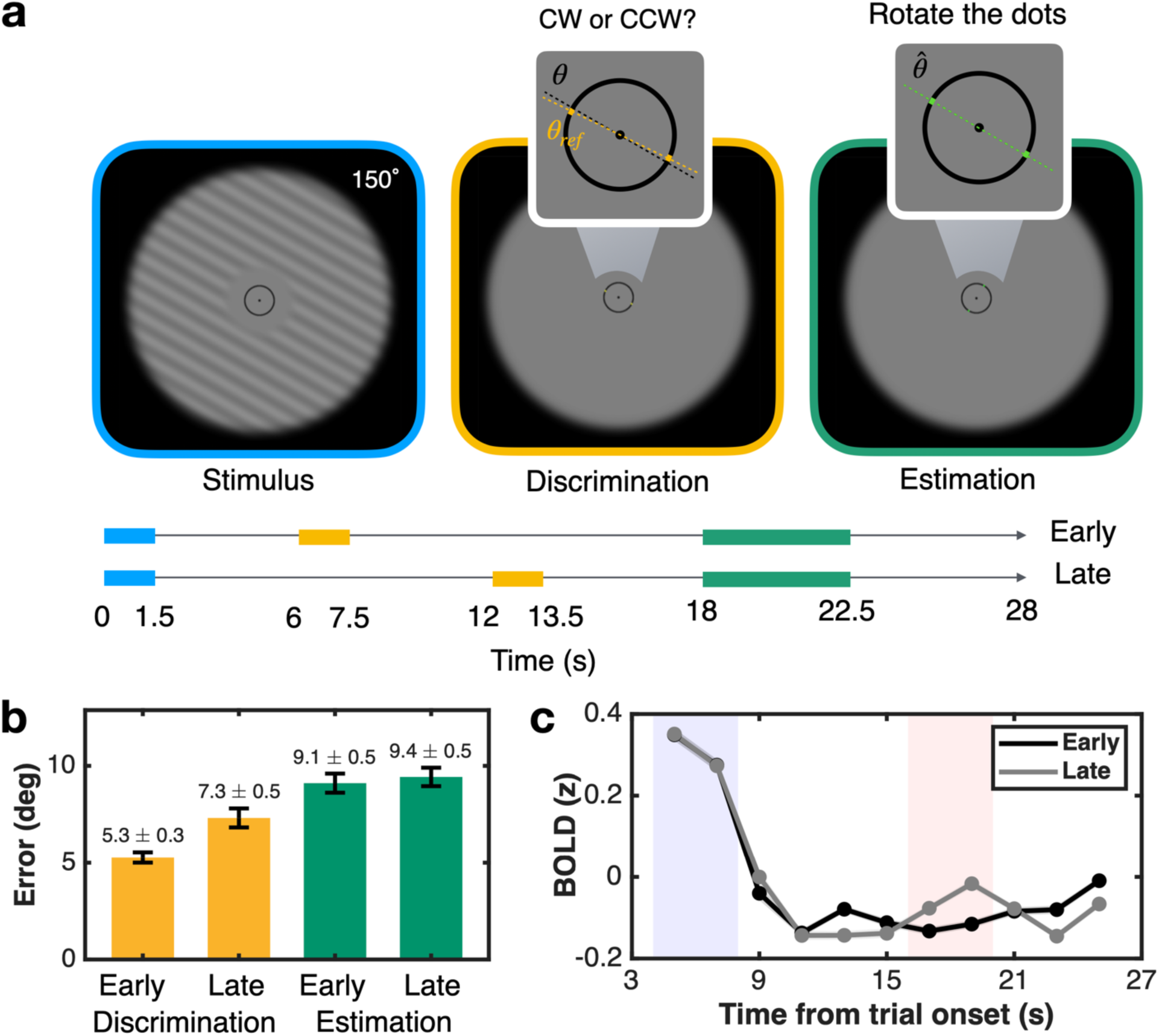
| Experimental paradigm, behavioral summary, and epoch definition. a, Delayed-estimation task used during fMRI. Top: Displays for the stimulus (blue outline), discrimination (yellow outline), and estimation (green outline) phases. Inset elements for the reference dots are colored for illustration only. Bottom: Task timeline showing the onset and duration of each phase relative to trial start. ‘Early’ and ‘Late’ denote the timing conditions for the discrimination phase. **b,** Behavioral performance. Mean error (in degrees) for the discrimination (yellow bars) and estimation (green bars) tasks, shown separately for early and late probe conditions. Error bars indicate s.e.m. across participants. Values above bars are mean ± s.e.m. across participants (N = 50). **c,** Definition of sensory and mnemonic epochs from EVC univariate BOLD time courses. Univariate z-scored BOLD responses for early (black) and late (grey) probe trials are plotted as a function of time from trial onset (symbols, group mean; shading, s.e.m.). Blue and red shaded regions indicate the sensory (4–8 s) and mnemonic (16–20 s) epochs, respectively.

## Results

### Temporal separation of sensory- and mnemonic-dominant epochs

To better separate late mnemonic activity from lingering stimulus-evoked responses—a central challenge in fMRI studies of WM—we designed a delayed-estimation task with an extended 16.5-s retention interval (**Fig. 2a**). Participants first viewed a single oriented grating (**Fig. 2a, blue**) and, after the prolonged delay, reported the remembered orientation (**Fig. 2a, green**). This extended delay allowed stimulus-locked BOLD responses to return closer to baseline, thereby making late-delay activity more likely to reflect mnemonic signals with minimal sensory contamination. To support continuous WM maintenance and probe potential interference, an oriented reference dot pair was presented either early or late during the delay, and participants judged the remembered target orientation relative to the reference (**Fig. 2a, yellow**).

Despite the demanding task structure and extended delay, participants exhibited robust memory performance. Mid-delay discrimination (**Fig. 2b, yellow bars**) and post-delay estimation (**Fig. 2b, green bars**) errors were both modest. Discrimination errors increased slightly from early to late probes and were smaller than post-delay estimation errors. This overall pattern, consistent with subtle degradation in mnemonic precision over time, aligns with prior findings that WM undergoes a process of diffusion or decay (**Baddeley, 1986**^26^**; Zhang & Luck, 2009**^27^**; Compte et al., 2000**^28^**; Wimmer et al., 2014**^29^**; Schneegans & Bays, 2018**^30^). Nevertheless, the overall low magnitude of errors across both tasks and throughout the prolonged delay indicates that orientation memory was reliably maintained, supporting the population-level analyses that follow.

To define distinct BOLD epochs dominated by sensory or mnemonic signals, we selected 4-s data windows that balanced signal robustness with minimal temporal overlap. The sensory epoch (4-s window centered at 6 s after stimulus onset) was chosen to capture peak stimulus-evoked activity, consistent with the canonical hemodynamic delay (**Fig. 2c, blue shade**). Crucially, the mnemonic epoch (4-s window immediately preceding the estimation period, centered at 18 s after stimulus onset) was selected because univariate BOLD responses had by that time returned to near-baseline levels following the initial stimulus-evoked activity (**Fig. 2c, red shade**). This temporal separation allowed us to isolate a late-delay epoch in which activity primarily reflected mnemonic content. To minimize potential contamination from the mid-delay probes as well, all subsequent multivariate analyses of late-delay activity within the mnemonic epoch were restricted to early-probe trials, where the reference appeared sufficiently early in the delay to prevent its residual sensory responses from affecting the defined mnemonic window. With these sensory and mnemonic epochs defined, we next examined their relationship using standard cross-decoding before turning to subspace and geometry directly.

### Cross-decoding generalizes poorly between sensory and mnemonic epochs

To situate our analyses relative to prior work, which largely relied on cross-decoding, we first examined the relationship between sensory and mnemonic representations within that framework. An inverted encoding model (IEM; **Brouwer & Heeger, 2009**^31^**; Sprague et al., 2016**^32^) was trained on multivoxel patterns from the sensory epoch (**Fig. 3a, blue shade**) and then tested across the full trial time course. The sensory-trained model effectively reconstructed orientation during the early sensory epoch. However, reconstruction fidelity declined markedly across the delay and reached baseline levels by the mnemonic epoch (**Fig. 3a, red shade**). Thus, a decoder trained on sensory-epoch activity generalized poorly to the late mnemonic epoch, particularly in early-probe trials where late-delay activity was least affected by probe-related contamination.

**Figure 3.**
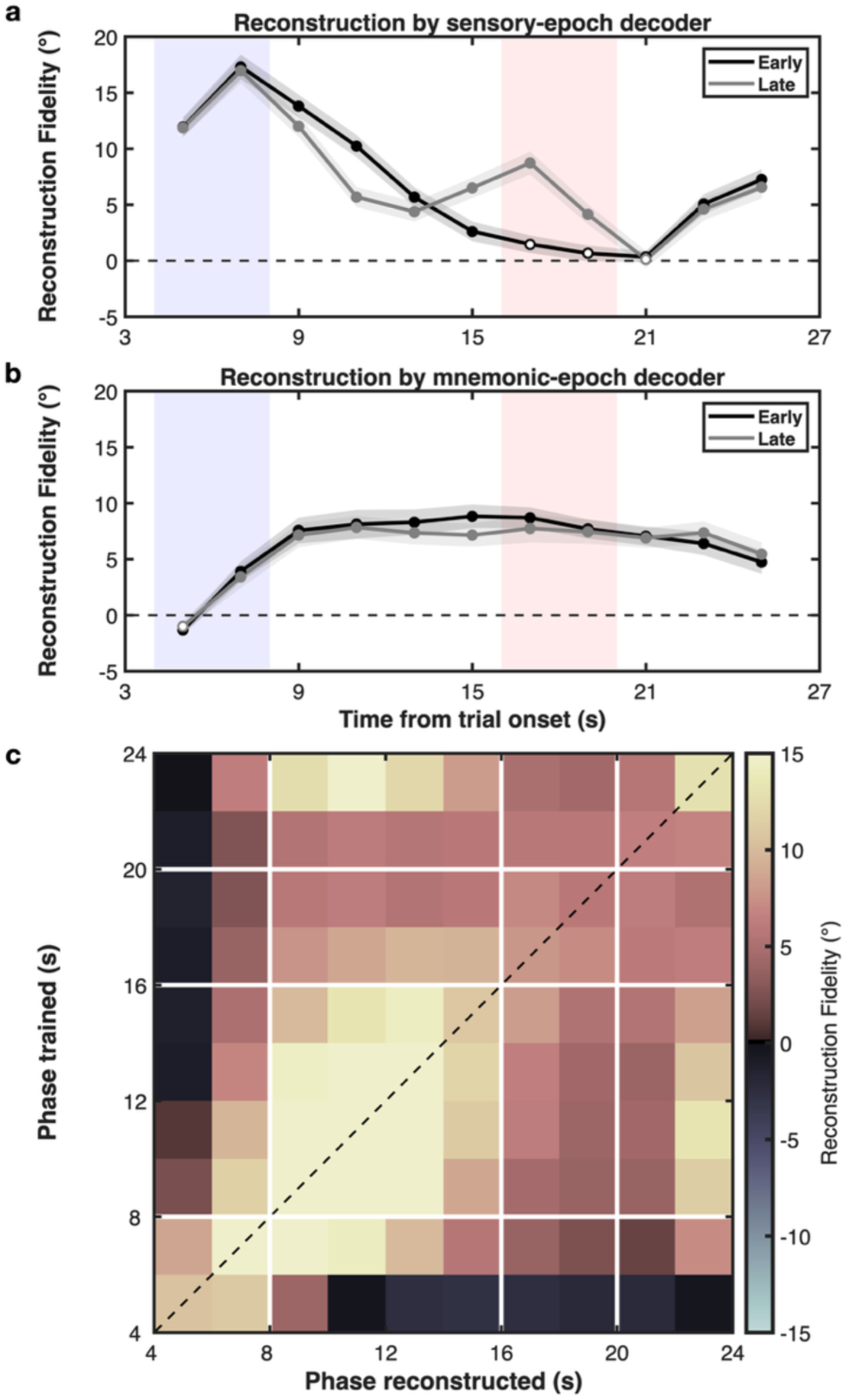
| Poor cross-epoch generalization in orientation decoding. a,b, Orientation reconstruction fidelity for the sensory-trained (a) and mnemonic-trained (b) decoders. Fidelity is plotted across time separately for early-probe (black) and late-probe (grey) trials (symbols, group mean; shading, s.e.m.). Open circles indicate time points at which reconstruction fidelity did not differ significantly from baseline. Blue and red translucent rectangles denote the sensory and mnemonic epochs, respectively. **c,** Time-resolved IEM reconstruction fidelity across all train–test time pairs for early-probe trials. Each cell shows reconstruction fidelity when the model is trained at one time point (y-axis) and tested at another (x-axis); the dashed diagonal indicates matched train and test times. Color indicates reconstruction fidelity.

Conversely, training the IEM on mnemonic-epoch activity from early-probe trials only (**Fig. 3b, red shade**) yielded robust orientation reconstructions during the late delay, confirming the presence of decodable mnemonic information in EVC. Yet this mnemonic-trained model generalized poorly to the sensory epoch, producing weak or absent orientation reconstructions there (**Fig. 3b, blue shade**). A time-resolved generalization analysis (**Fig. 3c**) further supported this pattern: temporal generalization matrices showed strong within-epoch decodability along the main diagonal of the sensory and mnemonic windows, but minimal generalization between them.

Thus, although orientation was reliably decodable from EVC during both the sensory and mnemonic epochs, decoders trained in one epoch generalized poorly to the other. This cross-decoding pattern indicates a substantial change in the mapping between orientation and multivoxel activity across epochs. Critically, however, as noted in the Introduction, cross-decoding failure does not by itself distinguish a distinct-code scenario from a geometry-preserving re-embedded-code scenario.

### Sensory and mnemonic orientation representations occupy separable low-dimensional subspaces

We therefore next examined the structural relationship between sensory and mnemonic representations in population activity space, with a focus on their low-dimensional organization. To isolate orientation-related structure while down-weighting shared task-locked dynamics, we applied demixed principal component analysis (dPCA; **Kobak et al., 2016**). dPCA separated orientation-related variance from condition-independent temporal variance and captured nearly as much variance as standard PCA (95.7% of the variance captured by the first 10 PCA components; **Supplementary Fig. 2b**), indicating that a small number of demixed components accounted for most of the population variance. For each participant, we defined a common low-dimensional state space from the three demixed components that explained the most orientation-dependent variance across the sensory and mnemonic epochs, and projected both epochs into this space. Results were qualitatively similar when more than three orientation-related dimensions were used (**Supplementary Fig. 2d**).

Within this common space, we defined an orientation subspace for each epoch as the two-dimensional plane that best fit the eight orientation-bin centroids (**Methods**). In both epochs, orientations traced a smooth, approximately circular polygon within the fitted plane (**Fig. 4a**), indicating ring-like orientation structure whose first two dimensions captured most of the orientation-dependent variance. To quantify separation between epochs, we measured the angle between the normals of the sensory and mnemonic planes in the 3D dPCA space. Across participants, this angle was large (mean = 71.26° ± 2.36° s.e.m.), substantially exceeding within-epoch plane-angle baselines estimated by bootstrap resampling (*p* < 10^−6^; **Fig. 4b**). At the individual level, 40 of 50 participants showed sensory-mnemonic plane angles significantly larger than their participant-specific common-subspace null distributions (**Fig. 4c**).

**Figure 4.**
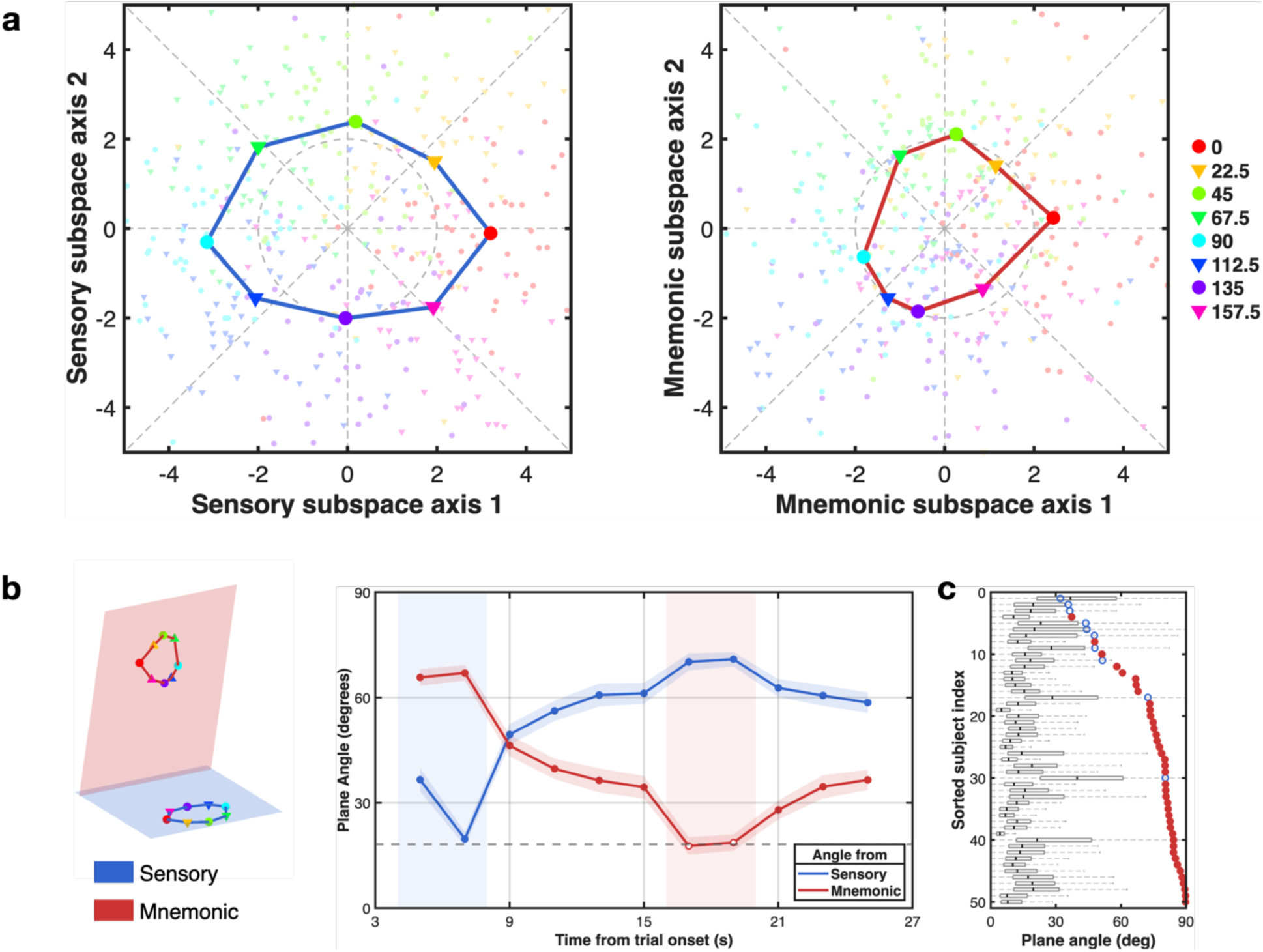
| Sensory and mnemonic orientation representations occupy separable low-dimensional dPCA subspaces. a, Orientation states during the sensory (left) and mnemonic (right) epochs projected into their corresponding fitted two-dimensional dPCA subspaces. Small symbols indicate orientation centroids for individual participants; large symbols show across-participant means. For visualization of group means, participant-specific coordinates were rotated within the corresponding subspace to a common orientation before averaging (**Methods**). Colors denote the eight circular orientation bins (see legend), revealing approximately ring-like structure in both epochs. b, Time-resolved alignment of the instantaneous orientation plane with the sensory reference plane (blue) and mnemonic reference plane (red), plotted as a function of time from trial onset. Using a fixed dPCA projection defined from the sensory and mnemonic epochs, we fitted an orientation plane at each time point and computed its angle relative to the sensory and mnemonic reference planes. The dashed line indicates the mean within-epoch benchmark from bootstrap resampling; shaded bands indicate mean ± s.e.m. across participants (N = 50). c, Observed sensory-mnemonic plane angles for individual participants overlaid on participant-specific common-subspace null distributions (*null*_0_; **Methods**). Filled markers indicate participants whose observed plane angle exceeded their null distribution at p < 0.05.

Time-resolved analyses using the same fixed dPCA projection further showed that the instantaneous orientation plane aligned more closely with the sensory reference subspace early in the trial and with the mnemonic reference subspace late in the delay (**Fig. 4b**). Intermediate time bins showed a gradual shift in alignment, which we interpret cautiously given the temporal blurring inherent to BOLD signals. Thus, these analyses indicate that sensory and mnemonic orientation representations in EVC occupy separable subspaces.

### Representational geometry is preserved across sensory and mnemonic epochs

Although dPCA is useful for isolating orientation-related subspaces and characterizing their spatial arrangement, it does not by itself establish whether representational geometry is preserved across epochs because it provides only a low-rank approximation of the full multivoxel state space. We therefore next quantified representational geometry directly from the native multivoxel patterns using cross-validated Mahalanobis (crossnobis) distance, which captures the pairwise distance structure among orientation-dependent activity patterns without relying on a low-dimensional approximation (**Diedrichsen et al., 2016**^33^**; Walther et al., 2016**^34^; see **Methods**).

Crossnobis-based representational dissimilarity matrices confirmed similarly structured orientation geometry across epochs (**Fig. 5a,d**). Within both the sensory and mnemonic epochs, crossnobis distances increased monotonically with circular orientation disparity, but not in an arbitrary way: the distance-disparity functions were well described by the circular chord-length rule expected for points arranged around a ring, *d*(Δ*θ*) = 2*R*sin (Δ*θ*/2) (**Fig. 5c, blue and red symbols**). In both epochs, distances increased steeply for nearby orientations and then progressively flattened for more distant orientation pairs, consistent with the nonlinear geometry of a ring-like representation rather than a purely linear ordering, as reflected in the close agreement between the empirical data and the chord-length fits in **Fig. 5c**. Correspondingly, sensory and mnemonic representational dissimilarity matrices showed similar structure, and their off-diagonal entries were reliably correlated across participants (mean Spearman’s ρ = 0.098 ± 0.020; p = 1.17 × 10⁻⁵), indicating that relative distances among orientations were preserved across epochs. Although the mean correlation may seem modest in magnitude, it was estimated separately within participants over the off-diagonal structure of 24 orientation conditions and consistently positive across participants, yielding a highly reliable effect. By contrast, cross-epoch dissimilarities between sensory and mnemonic patterns showed little dependence on orientation disparity (**Fig. 5b**) and a near-flat distance–disparity function (**Fig. 5c, grey dots**), in contrast to the structured within-epoch functions, consistent with the subspace separation identified by dPCA.

**Figure 5.**
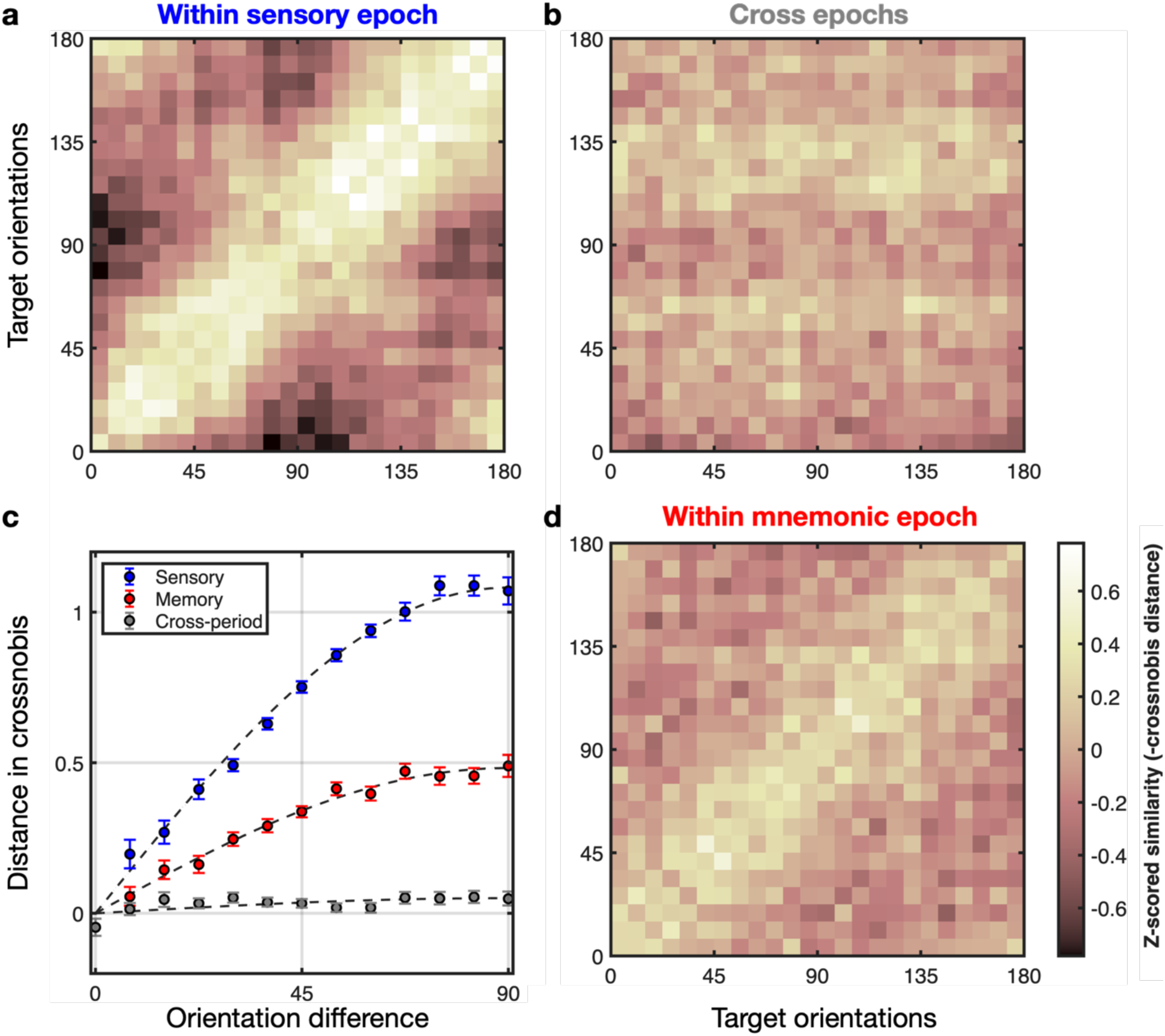
| Sensory and mnemonic orientation representations preserve representational geometry across epochs. a,d, Within-epoch representational dissimilarity matrices for the sensory (**a**) and mnemonic (**d**) epochs, averaged across participants (N = 50). For visualization, each cell shows participant-wise z-scored negative cross-validated Mahalanobis (crossnobis) distance between multivoxel patterns evoked by a pair of orientations within the same epoch; warmer colors indicate greater similarity. **b,** Cross-epoch matrix computed from crossnobis distances between sensory-epoch and mnemonic-epoch patterns for all orientation pairs. **c,** Mean crossnobis distance as a function of circular orientation disparity Δ*θ*for sensory (blue), mnemonic (red), and cross-epoch (gray) comparisons. For each participant, distances were averaged over all orientation pairs with the same Δ*θ*and then averaged across participants (symbols, mean ± s.e.m.). Dashed lines show fits of the circular chord-length function *d*(Δ*θ*) = 2*R*sin (Δ*θ*/2), with best-fit radii *R*_sen_ = 0.541, *R*_mne_ = 0.241, and *R*_cross_ = 0.025.

Together with the dPCA results, these analyses indicate that sensory and mnemonic orientation representations in EVC occupy separable subspaces while preserving a common ring-like organization of orientation space across epochs. This preserved geometry is most evident in the disparity-dependent distance structure expected for orientations arranged on a circle. We therefore interpret the sensory–mnemonic relationship as a re-embedded code: the two representations are expressed in separate subspaces while remaining geometrically related.

### Concurrent sensory and mnemonic readout during discrimination and estimation

Having shown that sensory and mnemonic orientation representations in EVC occupy separable subspaces while preserving representational geometry, we next asked whether these two information sources could be dissociated during task periods in which both were concurrently relevant. During the discrimination and estimation phases, participants viewed reference dot pairs while maintaining the target orientation in memory, creating epochs in which current sensory input and mnemonic information were both present. We therefore tested whether sensory- and mnemonic-trained decoders yielded dissociable readouts of these two information sources.

We first examined the mid-delay discrimination period (**Fig. 2a, yellow**), in which an oriented dot pair was briefly presented as a static reference and participants judged whether the remembered target was clockwise or counterclockwise relative to it (**Fig. 6a, top**). Thus, the discrimination period required EVC to represent a newly presented sensory reference while concurrently maintaining the target orientation in memory. For each trial, we applied the sensory-trained IEM and examined the reconstructed orientation (relative to the true target) as a function of reference offset (−21°, −4°, 0°, +4°, +21°). The reconstructed orientation showed a strong positive dependence on the reference (group-level regression slope from subject means = 0.939; test vs memory-aligned horizontal, slope = 0: t(49) = 6.07, p = 1.80 × 10^−+^, BF10 = 8.21 × 10^4^; test vs reference-aligned diagonal, slope = 1: t(49) = −0.39, p = 0.697, BF01 = 6.04; **Fig. 6a, middle**), indicating that the sensory-trained IEM primarily tracked the currently presented reference orientation. In contrast, when we applied the mnemonic-trained IEM to the same data, reconstructed orientations were largely invariant to the reference offset (slope = 0.231; vs 0: t(49) = 1.79, p = 0.079, BF01 = 1.48; **Fig. 6a, bottom**), consistent with a target-centered mnemonic readout rather than a reference-driven sensory readout. Thus, during discrimination, the two IEMs yielded dissociable information from the same EVC measurements: a reference-dependent sensory readout and a reference-invariant mnemonic readout.

**Figure 6.**
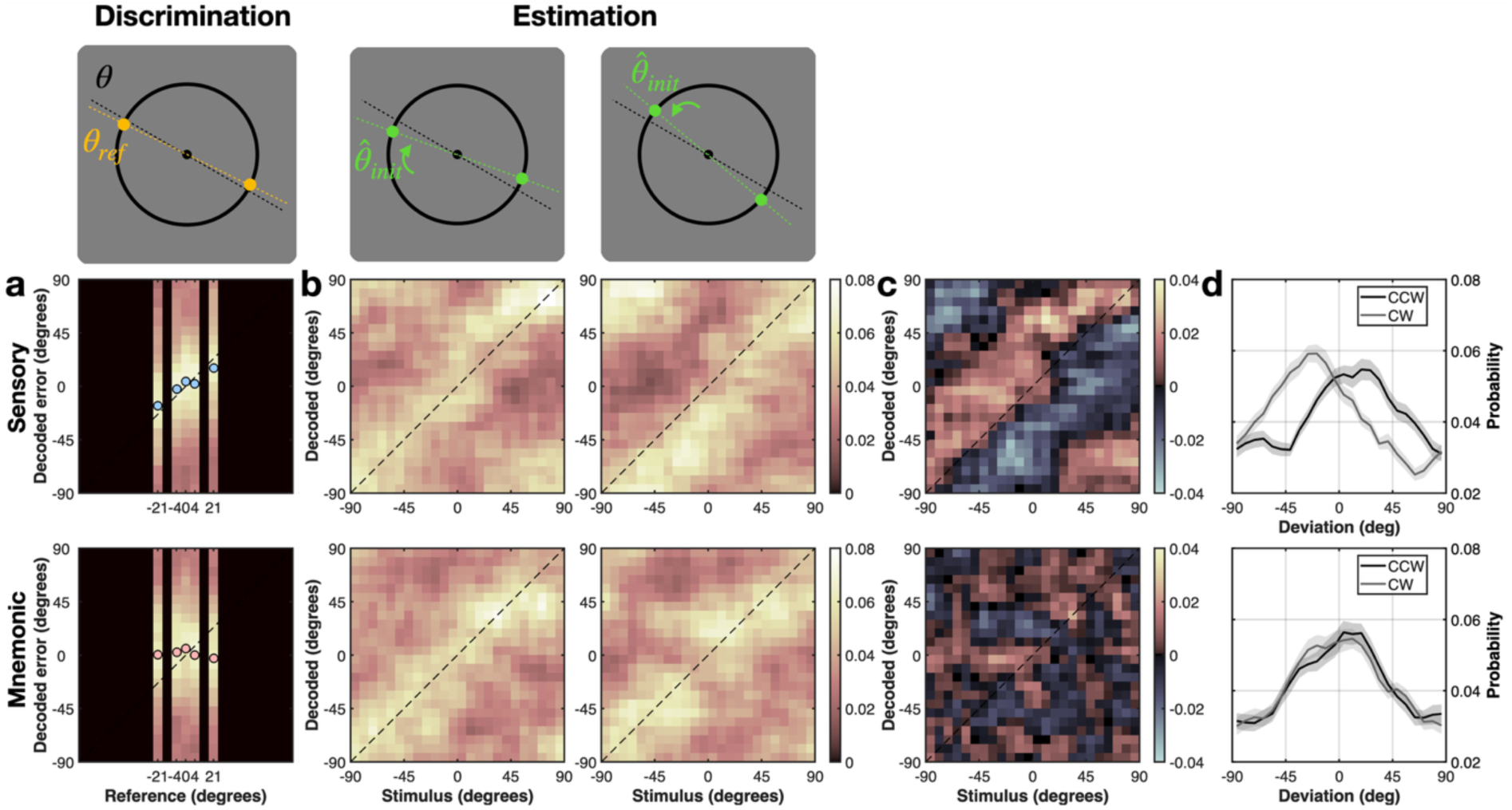
| Sensory- and mnemonic-trained IEMs dissociate concurrent sensory and mnemonic information during discrimination and estimation. a, Mid-delay discrimination. Top, task schematic: a remembered target orientation (*θ*) is compared to a briefly presented reference dot pair (*θ*_ref_). Middle and bottom, probability distributions of orientations reconstructed by the sensory-trained (middle row) and mnemonic-trained (bottom row) IEMs. For each reference-offset condition (−21°, −4°, 0°, +4°, +21° relative to the target), the reconstructed orientation distribution (relative to the true target) is shown. Colors indicate probability mass. Circles mark the expected decoded orientation for each reference offset, and diagonal dashed lines indicate the orientation aligned with the reference offset. **b–d,** Post-delay estimation. **b,** Top, task schematic: participants rotate the reference dot pair from an initial orientation (counterclockwise, CCW, or clockwise, CW, relative to the target; green arrows) to match the remembered target. Middle and bottom, probability distributions of orientations reconstructed by the sensory-trained (middle row) and mnemonic-trained (bottom row) IEMs. Reconstructed orientation probability is plotted as a function of target orientation, shown separately for trials in which the reference started on the CCW side (left column) versus the CW side (right column) of the target. **c,** Difference maps between CCW- and CW-conditioned distributions shown in **b** (CCW minus CW), highlighting start-side-dependent differences. **d,** Probability distributions of reconstructed orientation relative to the target, conditioned on the starting side of the reference (CCW vs CW) for the sensory-trained (top) and mnemonic-trained (bottom) IEMs, showing that sensory reconstructions depend on the starting side, whereas mnemonic reconstructions remain centered on the target.

We next asked whether a similar pattern was observed during the post-delay estimation period (**Fig. 2a, green**), when participants viewed a reference dot pair and rotated it from a starting orientation to match the remembered target (**Fig. 6b, top**). Trials were partitioned by whether the starting orientation of the reference lay counterclockwise (CCW) or clockwise (CW) relative to the target. Because the reference had to be rotated toward the target orientation, sensory input during estimation was concentrated along the segment between the reference’s starting orientation and the target orientation. Consistent with the discrimination-period results, reconstructions from the sensory-trained IEM were significantly biased toward the reference’s starting side (start-side bias in reconstructed error = 5.05°; t(49) = 6.65, p = 2.35 × 10^−-^, BF10 = 5.59 × 10⁵; equivalent CCW–CW difference in reconstructed orientation = 10.16°, t(49) = 6.75, p = 1.61 × 10^−-^; **Fig. 6b–d, sensory**). By contrast, reconstructions from the mnemonic-trained IEM remained centered on the remembered target and showed little dependence on the starting side (bias = 0.40°; t(49) = 0.53, p = 0.597, BF01 = 5.68; CCW–CW difference = 0.66°, t(49) = 0.44, p = 0.66; **Fig. 6b–d, mnemonic**).

Taken together, these analyses indicate that during both discrimination and estimation, sensory-and mnemonic-trained IEMs can dissociate concurrent sensory and mnemonic information from the same EVC activity patterns. This dissociation is consistent with the separable subspace organization identified above and extends it to task periods in which current sensory input and maintained mnemonic information coexist.

### Mnemonic orientation preferences are reorganized across voxels and decoupled from coarse retinotopic bias

A remaining question is whether mnemonic orientation coding preserves the voxel-wise preference structure and coarse retinotopic organization that characterize sensory encoding. If mnemonic activity were simply a reweighting or inversion of a fixed sensory map, voxel-wise preferred orientations should remain systematically related across epochs, and coarse retinotopic bias should be preserved. We therefore compared voxel-wise preferred orientations estimated separately in the sensory and mnemonic epochs and asked whether mnemonic coding retained the radial bias observed during sensory encoding.

When voxels were sorted by their sensory-epoch orientation preferences (see **Methods**), sensory responses showed a clear diagonal band (**Supplementary Fig. 4a**), consistent with an ordered orientation map, whereas applying the same ordering to mnemonic responses largely abolished that structure (**Supplementary Fig. 4b**). The reciprocal analysis produced the opposite pattern: sorting voxels by mnemonic preferences yielded a clear diagonal for mnemonic responses (**Supplementary Fig. 4d**), but not for sensory responses (**Supplementary Fig. 4c**). Quantitatively, participant-level circular correlations between voxel-wise sensory- and mnemonic-window preferred orientations were near zero (*r*_*circ*_ = -0.0041 ± 0.0122 s.e.m.; two-sided one-sample t-test on Fisher z-transformed participant correlations: t(49) = -0.33, p = 0.741), indicating that mnemonic-epoch orientation preferences were reorganized relative to sensory-epoch orientation preferences.

We next asked whether mnemonic coding retained the coarse retinotopic bias evident during sensory encoding, namely radial bias (**Freeman et al., 2011**^35^**, 2013**^36^**; Roth et al., 2018**^37^**; Ryu & Lee, 2024**^38^). To visualize the spatial organization of preferred orientation, we projected group-average preferred-orientation estimates onto cortical flatmaps of V1–V3 for the sensory and mnemonic epochs (**Fig. 7**). During the sensory epoch, preferred orientations exhibited a broad and smoothly varying large-scale organization consistent with coarse radial topography. During the mnemonic epoch, this large-scale organization was visibly redistributed and did not align with the radial-like structure observed during sensory encoding.

**Figure 7.**
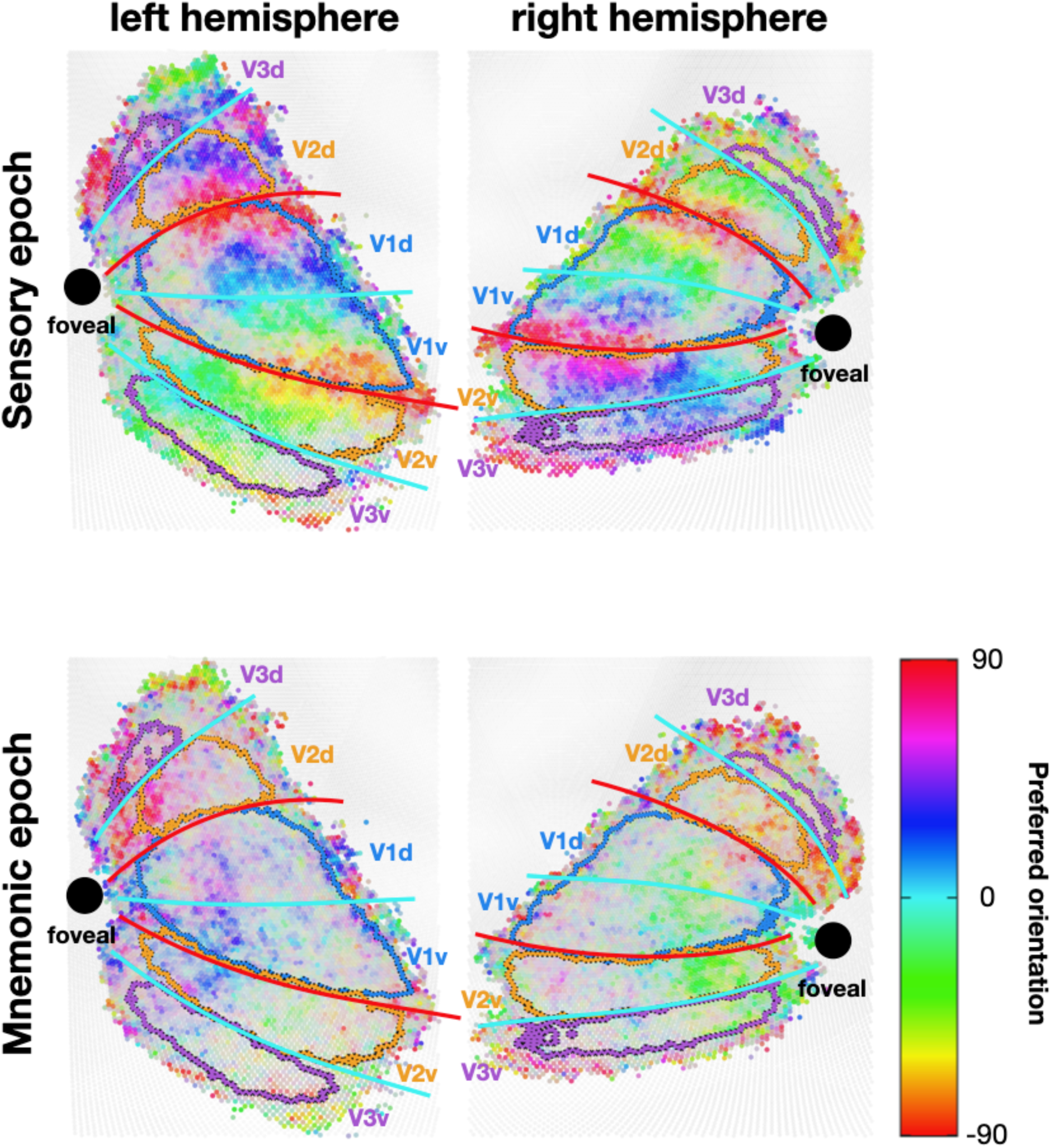
| Preferred-orientation maps in cortical space differ between sensory and mnemonic epochs. Group-average preferred-orientation maps projected onto fsaverage flatmaps for the left and right hemispheres. Top, sensory epoch. Bottom, mnemonic epoch. Colors indicate the circular-mean preferred orientation at each surface vertex. Overlaid contours delineate visual-area boundaries across early visual cortex. During the sensory epoch, preferred-orientation organization showed a broad large-scale structure consistent with coarse radial topography. During the mnemonic epoch, preferred-orientation organization appeared redistributed and did not align with the radial-like sensory structure observed during the sensory epoch.

Because **Fig. 7** provides a descriptive visualization across cortical space, we quantified radial bias separately in supplementary analyses. As we did not acquire full retinotopic maps, we used anatomy-derived retinotopic coordinates (**Benson et al., 2014**^39^) and restricted this analysis to V1, where template-derived visual-area assignments showed the strongest agreement with our independently defined visual-area labels (**Supplementary Fig. 3**). These analyses confirmed that the radial bias prominent in the sensory epoch was effectively absent in the late mnemonic epoch (**Supplementary Fig. 4f–g**). This lack of radial bias is unlikely to be explained by weaker late-delay reliability alone, because voxel-wise orientation preferences remained estimable during the mnemonic epoch even though they no longer aligned with radial retinotopic structure (**Supplementary Fig. 4a,d**).

Together, these results indicate that mnemonic orientation coding in EVC is reorganized relative to sensory coding and lacks the coarse retinotopic bias that characterizes sensory encoding. Thus, mnemonic orientation information can remain geometrically compatible with sensory coding without relying on coarse retinotopic bias.

### Mnemonic representations in EVC predict trial-to-trial behavior

So far, we have characterized the mnemonic representation in EVC in relation to sensory coding, focusing on its subspace organization and representational geometry. We next asked whether trial-by-trial variability in this mnemonic representation was related to behavioral variability across trials beyond that explained by the physical target orientation. This form of neural-behavioral linkage is often studied under the rubric of choice probability (**Britten et al., 1996**^40^**; Nienborg et al., 2012**^41^**; Choe et al., 2014**^42^**; Seidemann & Geisler, 2018**^43^**; Chicharro et al., 2021**^44^). Both the mid-delay discrimination and post-delay estimation tasks (**Fig. 2a**, yellow and green) require that the remembered target orientation remain accessible at behaviorally relevant moments during the trial. Consistent with prior work on postponed report (**Lemus et al., 2007**), if the mnemonic representation identified in EVC contributes to performance at those moments, then orientations reconstructed from delay-period BOLD activity should track trial-to-trial variability in discrimination choices and estimation reports at the relevant phases of the task.

To visualize this relationship during the discrimination phase, we first examined choice-conditioned reconstruction-error distributions (**Fig. 8a**). For both early- and late-probe trials, reconstruction errors from the mnemonic IEM showed systematic, though subtle, shifts between counterclockwise (CCW) and clockwise (CW) choices across reference-offset conditions, consistent with the modest effect sizes typically observed in choice-probability analyses of sensory cortex (**Haefner et al., 2013**^45^**; Choe et al., 2014**^42^**; Seidemann & Geisler, 2018**^43^**; Chicharro et al., 2021**^44^**; Lee et al., 2023**^46^).

**Figure 8.**
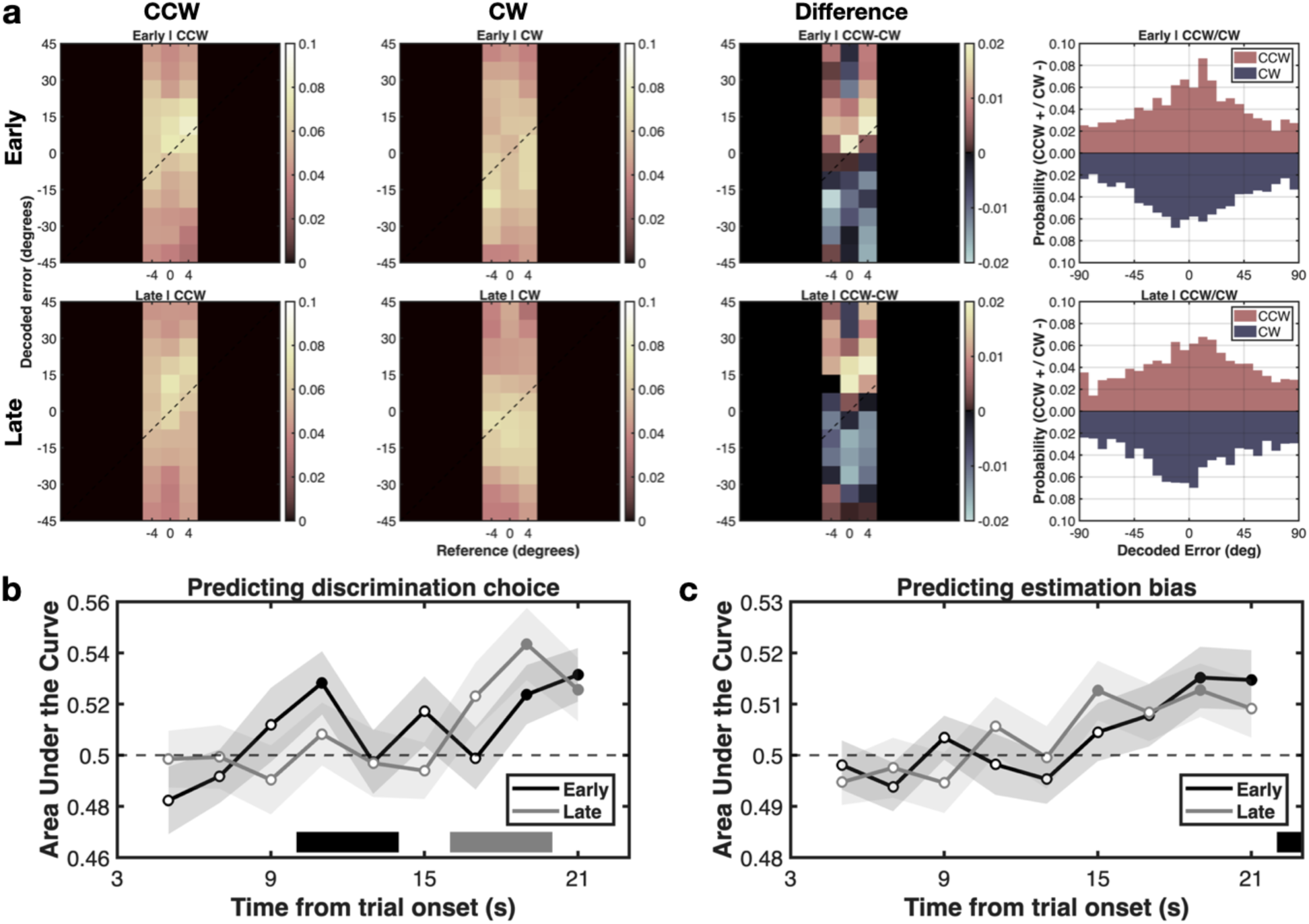
| Time-resolved prediction of discrimination choice and estimation bias from mnemonic reconstructions. a, Reconstruction-error distributions from the mnemonic-trained IEM during the discrimination phase, shown separately for early-probe (top row) and late-probe (bottom row) trials. For each reference offset (−4°, 0°, and +4°), distributions are split by discrimination choice (CCW vs CW). Difference panels show CCW − CW, and the rightmost histograms pool this choice difference after aligning errors to the reference across offsets. **b,** Time-resolved ROC AUC for predicting mid-delay discrimination choice from reconstructed orientations, shown separately for early-probe (black) and late-probe (grey) trials and averaged across reference offsets. c, Time-resolved ROC AUC for predicting estimation bias from reconstructed orientations, defined by median-splitting signed estimation error within each target orientation (CCW vs CW), again shown separately for early- and late-probe trials. **b,c,** lines show mean ± s.e.m. across participants; filled circles indicate time points significantly different from chance (0.5; two-sided one-sample t-test, p < 0.05, uncorrected). Horizontal bars denote the discrimination epochs in b and the estimation epoch in c.

We then quantified these effects over time. For each time point between 4 and 22 s after stimulus onset, we divided trials into two groups based on behavior and computed the receiver-operating-characteristic area under the curve (ROC AUC; chance = 0.5) for discriminating between the corresponding reconstructed-orientation distributions. First, reconstructed orientations were conditioned on binary mid-delay discrimination choices (clockwise vs counterclockwise relative to the reference), and time-resolved AUC was computed separately for early- and late-probe trials. Critically, the timing of choice predictability depended on when the discrimination judgment occurred. In early-probe trials, AUC rose above chance predominantly in an earlier window, time-locked (after accounting for the hemodynamic delay) to the early discrimination phase (**Fig. 8b**, black). In late-probe trials, the increase was shifted later in the trial, aligned with the late discrimination phase (**Fig. 8b**, grey). Thus, trial-by-trial variability in discrimination choice was reflected in the mnemonic representation at the behaviorally relevant time for each probe condition.

We next asked whether the same mnemonic representation also tracked biases in the final continuous estimation reports. Here, trials were grouped according to whether the behavioral estimate on that trial fell clockwise or counterclockwise relative to the median of the orientation-conditioned estimate distribution, and we again computed time-resolved AUC for the reconstructed orientations. In contrast to the probe-locked pattern observed for discrimination, estimation-related predictability was not tied to probe timing. Instead, it gradually increased over the course of the trial and peaked toward the end, closely tracking the post-delay estimation phase and doing so similarly for early- and late-probe trials (**Fig. 8c**). Thus, the mnemonic representation tracked trial-by-trial variability in the final continuous report specifically during the estimation phase. This pattern is consistent with a mnemonic representation that increasingly reflects the remembered orientation that ultimately guides the continuous report.

Together, these results show that trial-by-trial variability in mnemonic representations in EVC tracks both mid-delay discrimination choices and post-delay estimation reports at the behaviorally relevant stages of the trial. Thus, the mnemonic representation identified in EVC is linked not only to the remembered target orientation itself, but also to task-relevant variability in WM-guided judgments.

## DISCUSSION

### Population-level relationship between sensory and mnemonic representations in EVC

A central question for sensory recruitment accounts of WM is how EVC represents remembered information while continuing to process ongoing sensory input. Here, we addressed this question at the level of population representations. Using an fMRI paradigm that temporally separates sensory- and mnemonic-dominant epochs, together with analyses of decoding, low-dimensional subspace structure, and representational geometry, we asked how sensory and mnemonic orientation representations in human EVC are related.

Our results indicate that sensory and mnemonic orientation representations in EVC are neither fully shared nor entirely incompatible, but instead occupy separable subspaces while preserving representational geometry across epochs. This organization explains both the failure of cross-decoding between epochs and the dissociable readouts of concurrent sensory and mnemonic information, while remaining consistent with the observed reorganization of voxelwise orientation tuning and the lack of coarse retinotopic bias during the mnemonic epoch. Trial-by-trial variability in the mnemonic representation further predicted behavior across tasks.

Taken together, these findings support a population-level account in which EVC represents current sensory input and remembered orientation in related but dissociable formats. Rather than favoring either a single shared code or fully distinct and incompatible codes, our data are most consistent with a re-embedded-code relationship: sensory and mnemonic representations are expressed in separate subspaces while preserving their representational geometry. In this sense, EVC may support an “extended present” in which information from the immediate past and the current moment remains simultaneously available for behavior (**James, 1890**^1^).

### Separable subspaces with preserved geometry reconcile sensory recruitment with concurrent sensory processing

Our findings provide a population-level account of how sensory recruitment can be reconciled with concurrent sensory processing in EVC. A central challenge for this framework is how remembered information can be maintained without interfering with incoming sensory input when both are represented within the same cortical population (catastrophic interference; **Mendoza-Halliday et al., 2014**^9^**; Stokes, 2015**^10^**; Bettencourt & Xu, 2016**^11^**; Xu, 2017**^12^**, 2020**^13^). The present results suggest a specific representational organization through which this may be possible: sensory and mnemonic orientation signals occupy separable low-dimensional subspaces while preserving representational geometry across epochs. In this arrangement, subspace separation can, in principle, reduce interference between past and present information, whereas preserved geometry maintains a common representational format that supports their comparison or combination when behavior requires it.

This interpretation is further supported by the finding that sensory- and mnemonic-trained IEMs yielded dissociable readouts even when current sensory input and remembered orientation were concurrently relevant. Together with the behavioral linkage of the mnemonic representation, this suggests that late-delay coding in EVC is not merely a residual trace of earlier sensory activity, but is related to remembered information that remains available for WM-guided judgments. At the same time, because these inferences are based on fMRI population patterns, they should be interpreted at the level of representational organization rather than as direct evidence for a specific circuit mechanism.

More broadly, the present findings suggest that sensory recruitment should not be understood as requiring a single shared representational format between sensation (or perception) and memory. Instead, sensory recruitment in EVC may take the form of a re-embedded representational relationship, in which mnemonic information remains geometrically related to sensory coding while being expressed in a separable population subspace. This view preserves the core intuition of sensory recruitment—that remembered content is represented within sensory cortex—while helping explain how remembered information and current sensory input can remain concurrently available with limited interference.

More generally, a representational organization in which related information occupies separable subspaces while preserving internal geometry could provide a generic way for cortex to keep internally generated and externally driven signals simultaneously available without collapsing them into a single representational format. A related logic has been described in motor cortex, where preparatory activity is segregated from movement execution via null or orthogonal subspaces (**Kaufman et al., 2014**^21^). In this sense, the present results may point to a broader population-level principle for coordinating past and present information within a shared cortical region.

### Re-evaluating fMRI evidence for WM codes in EVC

The representational framework developed above helps clarify several seemingly conflicting observations from prior fMRI studies of WM in EVC. Classic studies have often been interpreted as evidence that mnemonic representations reuse the same feature-selective code that supports sensory processing, based largely on successful cross-decoding from stimulus-evoked responses to nominal delay-period activity (**Harrison & Tong, 2009**^3^**; Serences et al., 2009**^4^**; Ester et al., 2013**^47^). From the present perspective, however, cross-decoding alone cannot distinguish a fully shared code from a relationship in which sensory and mnemonic representations preserve geometry while being expressed in separable subspaces. Moreover, when delay periods are short, apparent cross-epoch generalization may also reflect residual stimulus-evoked activity within the hemodynamic response rather than a temporally isolated mnemonic representation.

The findings of **Rademaker et al. (2019)**^48^ are informative in this respect. They showed that remembered orientations could be reconstructed in the presence of sensory distractors using a decoder trained on stimulus-evoked responses. Yet the activity patterns used for reconstruction still fell within the temporal range of the target hemodynamic response, raising the possibility that the measured signals reflected a mixture of lingering stimulus-evoked and mnemonic activity. Under such conditions, a sensory-trained decoder would be expected to recover both current sensory input and residual target-related signals, consistent with the idea that concurrent sensory and mnemonic information can coexist in the same measured EVC patterns without necessarily sharing a single representational code.

A related line of work by Curtis and colleagues (**Kwak & Curtis, 2022**^49^**; Duan & Curtis, 2024**^50^) reported delay-period activity in EVC that aligned in a line-like fashion with remembered orientation and extended across retinotopic space, which they interpreted as evidence for abstract, priority-like mnemonic representations. Viewed from the present perspective, a key issue is whether that structure reflects temporally isolated mnemonic coding or instead remains shaped by lingering sensory organization. Because the short delay windows used in such studies overlap with the peak and early decay of the sensory hemodynamic response, mnemonic structure is difficult to separate from residual sensory activity that may still carry coarse retinotopic bias. This raises the possibility that at least part of the previously observed line-like structure reflects residual sensory topography rather than the structure of late mnemonic coding per se.

Taken together, prior fMRI work on WM in EVC spans a spectrum from shared-code interpretations to evidence for partially transformed or abstracted representations. The present framework suggests that some of this variation may arise from differences in temporal isolation and in the level of representational analysis.

### Possible neural bases for re-embedded mnemonic representations

Although our data are based on fMRI population patterns and therefore do not identify the underlying circuit mechanism, they do constrain plausible neural interpretations. In particular, the late-delay mnemonic representation does not look like a simple replay of the sensory response. Instead, it appears to preserve feature structure while being reorganized relative to sensory code and less tied to the coarse retinotopic organization that characterizes stimulus-driven activity.

One possible explanation is that mnemonic representations in EVC are shaped by feedback from higher-tier cortical areas. Along the visual hierarchy, receptive fields typically become larger and less tightly linked to precise retinal location, yielding more abstract and less retinotopically specific feature representations (**Sreenivasan et al., 2014**^51^**; Summerfield et al., 2020**^52^). If such higher-tier representations are fed back into EVC during the delay period, they could preserve orientation structure while reducing dependence on the coarse retinotopic bias that dominates stimulus-driven responses (**Freeman et al., 2011**^35^**, 2013**^36^**; Roth et al., 2018**^37^**; Ryu & Lee, 2024**^38^). On this view, mnemonic information would remain related to sensory coding while being re-expressed in a format that is less constrained by the original sensory topography.

A related possibility is that this shift reflects differential engagement of layer-specific neural populations within EVC. Anatomically, feedforward and feedback projections target different laminar compartments: feedforward input predominantly targets layer 4, whereas feedback projections preferentially engage superficial and deep layers while largely bypassing layer 4 (**Felleman & Van Essen, 1991**^53^**; Markov et al., 2014**^54^). Because an fMRI voxel pools activity across multiple layers, changes in the relative contribution of these subpopulations could alter the aggregate tuning profile measured from the same voxel. Under this interpretation, the observed voxel-level reorganization could arise because delay-period activity reflects a different mixture of sensory- and feedback-dominated populations than does stimulus-driven activity.

More broadly, these possibilities are consistent with known properties of higher-tier associative cortex, including mixed selectivity and high-dimensional population coding, which could support multiple related representational manifolds across distinct axes of activity (**Rigotti et al., 2013**^55^**; Fusi et al., 2016**^56^**; Bernardi et al., 2020**^57^). Recurrent network models likewise show that related information can be reorganized across partially distinct subspaces while preserving task-relevant structure (**Machens et al., 2010**^58^**; Mante et al., 2013**^59^). We do not claim that the present data demonstrate any one of these mechanisms. Rather, they suggest that re-embedded mnemonic representations in EVC may arise from interactions between stimulus-driven sensory coding and feedback- or context-dependent population dynamics that are less tightly tied to sensory topography.

Distinguishing among these possibilities will require measurements with finer spatial and temporal resolution than fMRI alone can provide. In particular, laminar-resolved imaging, multiarea recordings, and causal perturbations will be important for determining whether re-embedded mnemonic representations in EVC reflect feedback from higher-tier areas, differential engagement of layer-specific populations, local recurrent reorganization, or some combination of these processes.

### Limitations and future directions

Our findings provide a population-level account of how sensory and mnemonic orientation representations are related in EVC, but they also leave important questions open.

First, our conclusions are based on fMRI multivoxel patterns, which provide an indirect measure of neural activity. Although these measurements constrain the representational organization of sensory and mnemonic coding in EVC, they do not identify the neuronal or circuit-level processes that generate it. The voxel-level reorganization observed here is compatible with several possibilities, including feedback from higher-tier areas, differential engagement of layer-specific populations, local recurrent reorganization, or some combination of these processes. Distinguishing among these alternatives will require measurements with finer spatial and temporal resolution, including laminar-resolved imaging, multiarea recordings, and causal perturbations.

Second, the present study focused on a single feature, orientation, in a single sensory region. Future work should test whether similar relationships between subspace separation and preserved representational structure arise for other visual features, such as motion or color, and in other cortical areas that support WM. Such work will help determine whether the re-embedded relationship identified here reflects a more general principle for coordinating remembered and currently available information.

More broadly, the present findings suggest that the key question for sensory recruitment accounts may not simply be where remembered information is represented, but how mnemonic and sensory representations remain related without becoming conflated. Addressing this question will require linking population-level measurements such as those used here to the neural processes that shape representational organization over time. Progress will likely depend on integrating high-resolution imaging, multiregional recordings, computational modeling, and causal perturbation to understand how related but dissociable representations are coordinated across cortical systems.

## Materials and methods

### Participants

A total of 50 healthy adults (30 females; age 19–32 years) with normal or corrected-to-normal vision participated in the study. All participants provided written informed consent in accordance with protocols approved by the Institutional Review Board of Seoul National University. Participants were naïve to the purpose of the study, reported no history of neurological or psychiatric illness, and met standard MRI safety criteria. Each participant completed at least two, and in most cases all three, scanning sessions of the main fMRI experiment.

### Experimental procedure and stimuli

Stimuli were generated in MGL and presented via an LCD projector at 60 Hz. The display consisted of a black background with a gray circular Gaussian aperture spanning the peripheral visual field, together with a central fixation dot and fixation ring. Participants completed 20 task runs of 12 trials each across three scanning sessions, and completed a 1-h practice session before scanning.

On each trial, participants viewed an oriented grating for 1.5 s (inner/outer radii, 2°/8.5°; spatial frequency, 1 cycle/degree; contrast reversal at 8/3 Hz) and maintained its orientation across a 16.5-s blank delay. During the delay, an interim two-alternative forced-choice (2AFC) discrimination probe was presented: a dot-pair reference, defined by two yellow nonius dots on the fixation ring, appeared for 1.5 s and participants judged whether the remembered orientation was counterclockwise (CCW) or clockwise (CW) relative to the reference under moderate time pressure. The discrimination probe occurred either early or late in the delay, beginning at 4.5 s or 10.5 s after stimulus offset, respectively (**Fig. 2a**).

At the end of the delay, the estimation report was cued by a change in the fixation marker. Participants adjusted a dot-pair report frame, defined by two green nonius dots, to reproduce the remembered orientation within a 4.5-s response window. The report frame appeared at an initial orientation (*θ*_*init*_), which served as a transient sensory input during the estimation epoch; analyses of estimation-phase sensory readouts stratified trials by the signed deviation of *θ*_*init*_ relative to the target orientation (CCW versus CW start side; **Fig. 6b-d**). Trials were followed by a 5.5-s inter-trial interval, yielding a total trial duration of 28 s.

Stimulus orientation (θ) was sampled uniformly from a discrete set spanning 0–172.5° (24 values at 7.5° increments; orientation treated as 180°-periodic). Reference orientation (*θ*_*ref*_) was defined relative to the target by adding a signed offset (*Δ*_*ref*_ ∈ { −21°, −4°, 0°, +4°, +21°}) to the target orientation (*θ*_*ref*_ = *θ* + *Δ*_*ref*_; modulo 180°). Thus, *Δ*_*ref*_ determined whether the target was CW or CCW relative to the reference. Even when the target and reference were identical, participants were not informed of this and still made the same choice judgment. Analyses that stratified trials by reference offset used these five signed offset levels.

### Behavioral analyses

#### Estimation error and bias conditioning

(**Fig. 2b**). Estimation error was defined as the 180°-periodic difference between the reported orientation and the target, wrapped to (−90°, 90°). To remove stimulus-dependent systematic biases when summarizing estimation performance (**Fig. 2b**), errors were conditioned on stimulus orientation by estimating the circular-mean error as a function of stimulus orientation (24 bins, 7.5°) and fitting this bias function with a 12-component von Mises derivative basis (κ = 6) using ridge regression (α = 1×10−10). The predicted bias was subtracted from each trial’s raw error and re-wrapped, and estimation performance was quantified as the root-mean-squared error (RMSE) of these bias-conditioned errors, computed separately for early- and late-probe trials.

#### Discrimination psychometric fits

For each participant, CCW/CW choices in the mid-delay discrimination task were fit with a probit psychometric function of the signed reference offset (Δref), including a lapse parameter (λ ≤ 0.30) and a stimulus-dependent bias term estimated from the estimation data above. Separate sensory-noise parameters and bias weights were estimated for early- vs late-probe trials, and model parameters were optimized by multistart constrained maximum-likelihood (20 initializations).

### MRI acquisition and preprocessing

MRI data were acquired on a Siemens 3T Tim Trio scanner with a 32-channel head coil. T1-weighted anatomical images were collected at 0.8-mm isotropic resolution (TR = 2.4 s, TI = 1.0 s, TE = 2.19 ms, flip angle = 8°). Functional data were acquired with T2*-weighted echo-planar imaging (EPI) (2.3-mm isotropic voxels; TR = 2.0 s; TE = 30 ms; flip angle = 77°).

Preprocessing used fMRIPrep (v20.2.0) with fieldmap-free distortion correction (–use-syn-sdc). For each voxel, time series were converted to percent-signal change, and nuisance regressors were removed in a single linear model including white-matter and cerebrospinal-fluid (CSF) signals, six motion regressors, and discrete cosine bases for frequencies below 0.008 Hz. No spatial smoothing was applied. Resulting time series were z-scored voxel-wise within run for subsequent multivariate analyses.

### ROI definition

Early visual cortex (EVC) ROIs comprising V1–V3 were defined in each participant using standard traveling-wave retinotopic mapping with two 15° wedge bowties aligned to the vertical and horizontal meridians. Voxel-wise signal-to-noise ratio (SNR) was estimated from an independent hemodynamic impulse response function (HIRF) localizer scan consisting of a whole-field checkerboard impulse (radius, 8°; 1/24 Hz), computed as the amplitude at the stimulus frequency divided by the mean amplitude of harmonics above the third. Voxels with SNR < 2 were excluded. For all main population-level analyses, V1–V3 were pooled across hemispheres and dorsal/ventral subdivisions into a single EVC ROI.

### fMRI data analysis overview

To test how sensory and mnemonic representations in EVC are related at the population level, we performed complementary analyses of cross-decoding, low-dimensional subspace structure, representational geometry, and voxel-level tuning organization across sensory and mnemonic epochs. Across participants, the dataset comprised 241.68 ± 14.95 trials per participant (mean ± SD), including 122.74 ± 8.90 early-probe trials and 118.94 ± 10.96 late-probe trials.

### Univariate EVC BOLD time courses

For **Fig. 2c**, we computed univariate EVC responses by averaging z-scored BOLD across all voxels in the EVC ROI for each trial and TR, aligned to trial onset. For each participant, we then averaged these time courses across trials separately for the early- and late-probe conditions. Group time courses were obtained by averaging participant-wise time courses and plotting mean ± s.e.m. across participants at 2-s resolution from 4 to 26 s after trial onset.

### Definition of sensory and mnemonic population pattern matrices for subspace and geometry analyses

The sensory-evoked and late-delay mnemonic population patterns in EVC were defined and constructed as follows. We constructed voxel × condition activity matrices *S* (sensory) and *M* (mnemonic) from non-overlapping time windows within the same subset of trials, providing well-separated inputs for all subsequent multivariate analyses (dPCA, RDMs, and IEMs). To minimize contamination from the slow hemodynamic response and from mid-delay decision-related activity, we restricted all direct sensory–mnemonic code comparisons to early-probe trials. Within this fixed trial subset, we contrasted temporally separated windows that were chosen to (i) capture peak stimulus-locked activity and (ii) isolate late mnemonic activity after the BOLD responses to both the initial stimulus and the early discrimination probe (reference dots) had returned near baseline. Volumes were assigned to an analysis window when their acquisition mid-time fell within that window. Late-probe discrimination trials were used in separate analyses (e.g., **Fig. 3a,b, Fig 8**), but not in the construction of S and M.

#### Sensory (*S*) activity matrix

To capture peak stimulus-driven responses, we defined a sensory analysis window from 4 to 8 s after trial onset, aligned to the onset of the oriented-grating target (**Fig. 2c, blue shading**). For each early-probe discrimination trial, we averaged the multivoxel EVC activity across all volumes whose acquisition mid-times fell within this 4–8 s window, yielding a single voxelwise pattern vector for that trial. Stacking these voxelwise vectors across trials produced a trial × voxel matrix *X*_*S*_(*nTrials* × *nVoxels*). To obtain condition-averaged patterns for subsequent analyses, we grouped trials by target orientation (either 8 circular bins or 24 discrete orientations, depending on the analysis) and averaged rows of (*X*_*S*_ within each condition, yielding the sensory activity matrix *S* (*nCond* × *nVoxels*).

#### Mnemonic (*M*) activity matrix

To capture late delay-period mnemonic representations, we defined a mnemonic analysis window from 16 to 20 s after trial onset, immediately preceding the estimation period (**Fig. 2c, red**). For each early-discrimination trial, we averaged multivoxel EVC activity across all volumes whose acquisition mid-times fell within this 16–20 s window, yielding a voxelwise pattern vector per trial. Stacking these vectors across trials produced a trial × voxel matrix *X*_*M*_ (*nTrials* × *nVoxels*). As for the sensory matrix, we then averaged rows of *X*_*M*_ within each orientation to obtain the mnemonic activity matrix and the corresponding condition-mean matrix *M*(*nCond* × *nVoxels*). Because fMRI BOLD responses lag underlying neural activity by ∼4–6 s (**Boynton et al., 1996**), the 16–20 s BOLD window primarily reflects neural activity approximately 10–16 s after trial onset. In early-probe discrimination trials, the discrimination probe (reference dots) occurs at 4.5 s after stimulus offset (i.e., 6 s after trial onset), so its hemodynamic response peaks well before the mnemonic window. Likewise, the mnemonic window ends before the hemodynamic response to the subsequent estimation report (triggered at 18 s) rises substantially. Thus, the 16–20 s epoch is maximally separated from both the initial stimulus-evoked response and the end-of-trial estimation-related response. Because *S* and *M* are defined from non-overlapping windows within the same early-probe discrimination trial subset, differences between them reflect temporal evolution of the representational code rather than differences in trial type or task demands.

### Multivariate pattern analysis: inverted encoding model (IEM) for sensory-mnemonic cross-decoding

We reconstructed trial-wise orientations from multivoxel BOLD patterns in EVC (pooled V1–V3) using an inverted encoding model (IEM; **Brouwer & Heeger, 2009**). This analysis was used both to quantify within-epoch orientation information and to test cross-epoch temporal generalization, asking whether an IEM trained on the stimulus-evoked sensory epoch generalized to the late-delay mnemonic epoch, and vice versa.

#### Linear encoding model

At each analysis time point (or time-averaged window), let *X* denote the multivoxel response matrix (*nTrials* × *nVoxels*) and let *C* denote the predicted channel-response matrix (*nTrials* × *nChannels*) obtained by evaluating the channel basis functions at each trial’s stimulus orientation. We assume a linear mixing model:

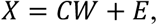

where *W* (*nChannels* × *nVoxels*) contains voxel-wise channel weights and *E* is residual noise. We estimated weights on the training set by ordinary least squares,

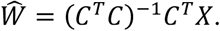

Given *Ŵ* channel responses for held-out test patterns *X_test_* were reconstructed by linear inversion:

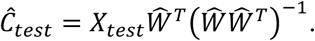

Weight estimation and reconstruction were performed independently within each cross-validation split using training data from the remaining runs only, thereby avoiding train-test leakage.

#### Cross-validation

All reconstructions were built on held-out data under leave-one-run-out cross-validation. Trials were grouped by run (12 trials per run). On each split, one run was held out for testing, and the remaining runs were used for training. This run-wise partitioning preserves local temporal structure and reduces inflation of reconstruction performance due to temporal autocorrelation.

#### Channel basis and dense reconstructions

Channel responses were defined over 180°-periodic orientation space using uniformly spaced cosine-power basis functions. For a channel centered at *ψ_k_*, the basis was

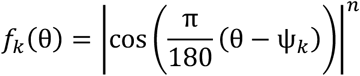

with *nChannels* = 8 and exponent *p* = 8. Because *p* is even, the basis is equivalent to |*cos*(·)|^*p*^, implementing 180°-periodic orientation tuning. To obtain smooth reconstructions, we repeated the model across phase offsets (1.5° shifts of the channel center) and concatenated the outputs to yield a dense reconstruction with 120 effective channel centers spanning 0–180°.

#### Training windows and temporal generalization

We trained two decoders that differed only in the time window used to define training patterns: a sensory-trained decoder using activity averaged over 4–8 s after trial onset (the sensory epoch), and a mnemonic-trained decoder using activity averaged over 16–20 s after trial onset (the mnemonic epoch). Each trained model was then applied to test patterns across time to obtain within-epoch decoding and cross-epoch generalization profiles (**Fig. 3a,b**). For the temporal generalization matrix (**Fig. 3c**), the same IEM procedure was repeated separately for each 2-s training bin, and each model was evaluated at all test time bins, yielding a train-by-test map of reconstruction fidelity.

#### Readout and decoding metric

For each trial and test time point, the reconstructed 120-point channel profile *c*^ was converted to a single decoded orientation using a doubled-angle population-vector readout,

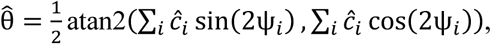

where *c*_*i*_ is the reconstructed response at channel center *ψ_i_*. Reconstruction error was defined as the 180°-periodic difference between 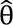 and the true orientation, wrapped to (−90°, 90°). Reconstruction fidelity was summarized as *fidelidy* = 45° − circular mean(|*error*|), using a doubled-angle circular mean. Under this definition, fidelity equals 0° at chance, because uniformly distributed errors yield a mean absolute error of 45°, and increases toward 45° as errors approach 0°.

#### Analyses of reconstructed orientation distributions during discrimination and estimation

The same trial-wise reconstructed orientations described above were used to characterize the outputs of the sensory-and mnemonic-trained IEMs during task periods in which current sensory input and maintained mnemonic information were concurrently relevant (**Fig. 6**).

#### Reference-conditioned reconstructed distributions during discrimination

For the mid-delay discrimination analysis (**Fig. 6a**), we analyzed decoded deviations separately for the five signed reference offsets (Δref = −21°, −4°, 0°, +4°, +21°) using the sensory-trained and mnemonic-trained IEMs applied to the discrimination-period test window. For visualization, decoded deviations were binned in 7.5° bins, each bin was pooled with its two adjacent bins with circular wrap-around at ±90°, and the pooled counts were renormalized to form a probability distribution for each reference offset. To obtain the point summaries used in **Fig. 6a**, we computed, within each participant and reference condition, a representative decoded deviation as the circular mean of the smoothed decoded-deviation distribution. We then regressed these five representative decoded deviations on signed reference offset, yielding one slope per participant and decoder. Group-level evidence for reference dependence was assessed with one-sample t-tests of participant-specific slopes against 0, corresponding to a memory-aligned readout, and against 1, corresponding to a reference-aligned readout. Bayes factors for these one-sample comparisons were computed using a Jeffreys-Zellner-Siow (JZS) Bayes t-test.

#### Start-frame-conditioned reconstructed distributions during estimation

For the post-delay estimation analyses (**Fig. 6b–d**), we analyzed decoded orientations from the sensory-trained and mnemonic-trained IEMs at the first time bin of the estimation epoch (22-24 s after trial onset). Trials were classified according to the starting orientation of the report frame relative to the target, defined by the sign of the wrapped difference between the starting frame orientation and the target orientation: counterclockwise (CCW) or clockwise (CW). For visualization in **Fig. 6b,c**, stimulus orientations were grouped into 24 bins (7.5° spacing), with each bin pooling the central orientation and its two neighboring orientations with wrap-around. Within each stimulus bin and start-side condition, decoded orientations were accumulated into 7.5° histograms, pooled with the two adjacent decoded-orientation bins using circular wrap-around, and renormalized. Separate CCW and CW decoded-orientation maps were constructed in this way, and the difference maps in **Fig. 6c** show CCW minus CW. For **Fig. 6d**, participant-specific decoded-deviation distributions were obtained separately for CCW and CW trials after collapsing across stimulus bins, normalized within participant, and then averaged across participants to yield mean ± s.e.m. curves.

#### Statistical summary of start-side effects

To quantify start-side bias, we computed, for each participant and IEM, the sign-aligned mean decoded deviation, defined as mean(sign(Δstart) × decoded deviation), where Δstart denotes the wrapped difference between the starting frame orientation and the target orientation. Positive values therefore indicate decoded shifts toward the starting side. Group-level evidence for start-side bias was assessed with a one-sample t-test against 0 across participants. To summarize the separation between the two start-side conditions, we also computed, for each participant and IEM, the difference between the mean decoded deviation on CCW versus CW trials after collapsing across stimulus bins, and tested this participant-level CCW–CW difference against 0 with a one-sample t-test.

### Demixed principal component analysis (dPCA) and sensory–mnemonic subspace of orientation codes

We used demixed principal component analysis (dPCA; Kobak et al., 2016) to construct a low-dimensional orientation state space that isolates stimulus-related variance while factoring out condition-independent BOLD dynamics, and to quantify the subspace relation between sensory and mnemonic orientation manifolds.

#### dPCA setup (conditions, factors, and time bins)

For each participant and ROI, we considered early-probe discrimination trials and grouped target orientations into 8 circular bins (22.5° spacing; each bin pooled three adjacent stimulus orientations with wrap-around). Within each orientation bin and each time bin, voxel patterns were averaged across trials to obtain condition means. These means defined an orientation × time × voxel data tensor.

We specified two task factors, orientation and time. The time-only marginalization captured condition-independent activity (shared temporal/BOLD dynamics), whereas the orientation main effect and the orientation × time interaction jointly defined stimulus-related activity (orientation structure that could vary across time). The time factor comprised four 2-s bins spanning the sensory (4–8 s; two bins) and mnemonic (16–20 s; two bins) windows.

dPCA decomposed the condition-averaged activity tensor into marginalizations corresponding to time-only, orientation-only, and orientation-by-time structure, and identifies paired decoder (W) and encoder (V) axes that capture these marginalized signals in a low-dimensional form. We focused on components assigned to the stimulus-related marginalizations (orientation and orientation × time), and used the resulting decoder matrix to project voxel activity into the low-dimensional orientation state space.

#### Regularization and component selection

For each participant, model selection proceeded in two stages. First, we performed a coarse cross-validated search over the total number of dPCA components (numComps = 10, 12, 15, or 20) and a broad logarithmically spaced grid of regularization values. For each candidate component count, dpca_optimizeLambda was run for 2 cross-validation repetitions, and the component-count/lambda pair with the lowest mean normalized reconstruction error was selected. Second, holding the selected component count fixed, we performed a fine lambda search over 25 values spanning one order of magnitude around the coarse optimum, using 100 cross-validation repetitions. We then fit the final dPCA model using the selected component count and lambda.

For all subsequent geometry analyses, we retained the leading three stimulus-related dPCA components to define a 3D orientation state space. These three components captured the large majority of orientation-dependent variance across the sensory and mnemonic epochs (**Supplementary Fig. 2**), while permitting 2D planar fits to the orientation manifold. The corresponding projection matrix, fitted on the four time bins spanning the sensory and mnemonic epochs, was reused for the time-resolved subspace analysis described below.

#### 2D visualization of sensory and mnemonic manifolds

For display, we computed eight orientation-bin centroids separately for the sensory and mnemonic epochs in the fixed 3D stimulus-related dPCA space. For each epoch separately, we performed PCA on the centered 8 × 3 point cloud of orientation-bin centroids and defined the epoch-specific 2D plane as the subspace spanned by the first two principal axes. Before averaging across participants, coordinates were centered and aligned within the corresponding plane by reflection/rotation to match the circular ordering of orientation bins across participants (**Fig. 4a**). These alignment steps were used for visualization only and did not affect any plane-angle calculations.

#### Plane definition and plane-angle metric

Using the same sensory and mnemonic centroid clouds, we quantified subspace separation as the angle between the unit normals of the two PCA-defined planes. Let ***n*_*S*_** and ***n*_*M*_** denote the unit normals of the sensory and mnemonic planes, respectively. Sensory–mnemonic subspace separation was quantified as the angle between these normals, computed as ***θ*** = ***arccos***(|***n*_*S*_** · ***n*_*M*_**|) and reported on a 0–90° scale. The absolute value treats planes with flipped normals as equivalent.

#### Time-resolved subspace alignment

To visualize how orientation geometry evolves over time phases in the fixed dPCA space (**Fig. 4b**), we computed orientation-bin centroids at successive 2-s time bins and, at each time bin, defined an instantaneous plane by PCA of the centered 8 × 3 centroid cloud. We then computed its angle relative to the sensory and mnemonic reference planes.

#### Statistical testing of subspace angle

For each participant, we estimated uncertainty on the sensory and mnemonic planes, and their separation angle, via bootstrap resampling of trials with replacement within each orientation bin (1,000 iterations). On each iteration, we recomputed orientation-bin centroids, fit epoch-specific planes in the fixed 3D stimulus-related dPCA space, and calculated the sensory–mnemonic plane angle. Participant-wise 95% confidence intervals were obtained from the 2.5th and 97.5th percentiles of the bootstrap distribution. To benchmark cross-epoch rotation against within-epoch variability, we also constructed participant-specific common-subspace null distributions (*null*_0_) by computing plane angles between independent bootstrap resamples drawn from the same epoch window (sensory-with-sensory and mnemonic-with-mnemonic) and pooling these within participants. These *null*_0_distributions reflect the angles expected if sensory and mnemonic data were drawn from a single underlying plane, and were used for the individual-level tests against a common-subspace hypothesis in **Fig. 4c**.

### Representational similarity analysis (RSA) of sensory and mnemonic orientation geometry

We used representational similarity analysis (RSA; Kriegeskorte et al., 2008) with cross-validated Mahalanobis (crossnobis) distances to quantify orientation-dependent similarity structure in multivoxel EVC patterns and to compare representational geometry between the sensory and mnemonic epochs.

#### Orientation binning and epoch patterns

Stimulus orientations were grouped into 24 bins (7.5° spacing), corresponding exactly to the discrete orientations in the stimulus set used in the experiment. For each trial, EVC multivoxel patterns were averaged within the sensory epoch (4–8 s) and within the mnemonic epoch (16–20 s) defined above, yielding one trial-wise pattern for each epoch. These trial-wise patterns served as inputs for constructing the sensory-epoch and mnemonic-epoch representational dissimilarity matrices (RDMs) (**Fig. 5a,d**). For the cross-epoch RDM, we used the same 24 orientation bins but computed dissimilarities between sensory-epoch and mnemonic-epoch patterns. Specifically, each matrix entry indexed the crossnobis distance between the sensory pattern for orientation i and the mnemonic pattern for orientation j, yielding a 24 × 24 matrix of between-epoch dissimilarities (**Fig. 5b**).

#### Cross-validated Mahalanobis (crossnobis) RDMs

For the sensory, mnemonic, and cross-epoch comparisons, we computed a 24 × 24 RDM using cross-validated squared Mahalanobis distance (“crossnobis” distance; **Diedrichsen et al., 2016; Walther et al., 2016**). We used 5-fold cross-validation stratified by orientation bin. Within each fold, trials in each orientation bin were partitioned into training and test sets, voxel patterns in both sets were mean-centered using the training-set voxel mean, and condition means were computed separately in the training and test partitions. Voxel-wise noise covariance was estimated from leave-one-out residuals (trial-wise pattern minus orientation-bin mean) within the training partition and regularized before distance estimation. Separate covariance estimates were used for the sensory and mnemonic within-epoch RDMs, whereas a pooled covariance estimate from sensory and mnemonic residuals was used for the cross-epoch RDM.

Let 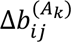 and 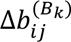 denote the noise-whitened voxel activity pattern (*b*) difference vectors between orientation *i* and j estimated from two independent partitions *A*_*k*_ and *B*_*k*_ in fold *k*. The crossnobis distance *d*_*i*j_ between orientations *i* and j was then computed as the dot product of the corresponding noise-whitened pattern difference vectors estimated on independent folds, and averaged across folds:

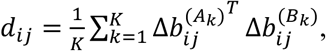

where *K* = 5 is the number of cross-validation folds. This quantity is an unbiased estimator of the true squared Mahalanobis distance and can take negative values when the underlying distance is small relative to noise. For the cross-epoch RDM, *d*_*i*j_was computed analogously between the sensory pattern for orientation i and the mnemonic pattern for orientation j using the pooled covariance estimate.

#### Group-level visualization of similarity matrices

For visualization in **Fig. 5a,b,d**, dissimilarities were converted to similarities as RSA = -*d*^2^. For the sensory and mnemonic within-epoch matrices, off-diagonal entries were z-scored within each participant before averaging across participants. For the cross-epoch matrix, all finite entries were z-scored within each participant before averaging. The resulting normalized matrices were then averaged across participants to obtain the group-level sensory, mnemonic, and cross-epoch representational similarity matrices shown in **Fig. 5a,b,d**.

#### RDM-similarity summaries (global geometry)

To quantify preservation of global representational structure across epochs, we compared the participant-specific sensory and mnemonic within-epoch matrices directly. For each participant, we vectorized the upper-triangular off-diagonal entries of the 24 × 24 sensory and mnemonic matrices and computed Spearman’s rank correlation (rho) between the two vectors. Because the within-epoch matrices are symmetric, only unique pairwise entries were used. Subject-level correlation values were summarized as mean ± s.e.m., and group-level evidence for positive correspondence was assessed with a one-sample t-test against zero across participants.

#### Distance-by-disparity summaries and chord-length fit

To summarize representational geometry as a function of circular orientation disparity (**Fig. 5c**), we converted the normalized similarity matrices back to z-scored crossnobis distances. Circular disparity between orientations i and j was defined as *Δθ*_*i*j_ = min(|*θ*_*i*_ − *θ*_j_|, 180° − |*θ*_*i*_ − *θ*_j_|). For the sensory and mnemonic matrices, distances were summarized over upper-triangular off-diagonal entries only; for the cross-epoch matrix, all finite sensory-mnemonic entries were included. Within each participant, distances were averaged across all orientation pairs sharing the same circular disparity, yielding one disparity-tuning curve for each matrix. These participant-specific curves were then averaged across participants and plotted as mean ± s.e.m. Dashed curves show least-squares fits of a chord-length model with free offset, *y* = *b* + 2*R* sin(*Δθ*), where b is an additive offset and R is the fitted radius. For display as relative distance, the fitted offset was subtracted from each curve before plotting. These fits were used as descriptive summaries of the disparity-dependent geometry rather than as separate subject-level inferential statistics.

### Voxel-wise orientation tuning, cortical-space preferred-orientation maps, and radial-bias analyses

To examine how orientation information was distributed across voxels in EVC, how this organization changed between sensory and mnemonic epochs, and how preferred orientation was arranged across cortical space (**Fig. 7**; **Supplementary Figs. 3–6**), we estimated voxel-wise orientation tuning from the pooled V1–V3 EVC ROI using the same early-probe trials used in the main sensory–mnemonic comparisons.

#### Voxel-wise orientation tuning model

We treated each voxel as a 180°-periodic orientation-tuned unit. For each participant, voxel, and time bin, trial-wise responses were modeled with a doubled-angle linear regression of the form

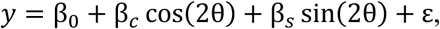

where y is the voxel response on a given trial, θ is the target orientation on that trial, and the doubled-angle terms capture the 180° periodicity of orientation. Models were fit by ordinary least squares using all early-probe trials. Preferred orientation and tuning strength were derived from the fitted coefficients as

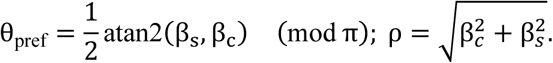

Thus, *θ*_*pref*_ is defined on [0, π), equivalent to [0°, 180°), and ρ indexes tuning strength. In addition to per-bin estimates, we also fit the model to trial-wise responses averaged within a sensory window and a mnemonic window, yielding time-averaged preferred-orientation estimates for the voxel-profile and cortical-space analyses described below. To provide a compact representation of both preference direction and selectivity, each voxel can be represented by a tuning vector

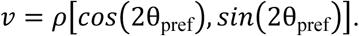

#### Voxel-profile matrices

To visualize how stimulus orientation was represented across voxels with different preferences, we constructed voxel-profile matrices (**Supplementary Fig. 4a–d**). For each participant, voxels were binned into 24 preferred-orientation bins (7.5° spacing) based on either sensory-window or mnemonic-window *θ*_*pref*_, depending on the analysis. For each stimulus orientation (24 bins, 7.5° spacing) and each preference bin, we then averaged BOLD responses across trials of that stimulus orientation and across voxels in the bin, using responses from the sensory window or mnemonic window as appropriate. This yielded a 24 × 24 stimulus-by-preference matrix per participant and epoch, and participant-wise matrices were averaged for display.

#### Across-epoch preference correspondence

To quantify whether voxel-wise orientation preferences were preserved across epochs, we computed, for each participant, the circular-circular correlation between voxel preferred orientations estimated from the sensory and mnemonic windows across voxels with finite estimates in both windows. Because orientation preference is 180°-periodic, correlations were computed on doubled angles. Participant-level correlation coefficients were summarized as mean ± s.e.m. across participants, and group-level evidence for a non-zero across-epoch correspondence was assessed with a two-sided one-sample t-test against zero on Fisher z-transformed subject-level correlations. All participants exceeded the minimum valid-voxel threshold required for this analysis.

#### Cortical-space preferred-orientation maps

To visualize how preferred orientation was organized across cortical space (**Fig. 7**), voxel-wise preferred orientations from the sensory and mnemonic windows were projected from each participant’s full V1–V3 voxel mask to fsaverage separately for the left and right hemispheres. Subject-specific mid-gray surface vertices were transformed into EPI voxel space, matched to the empirical voxel coordinates, and resampled to fsaverage using nearest-neighbor correspondence on the spherical registration. At each fsaverage vertex, we computed the across-participant circular-mean preferred orientation separately for the sensory and mnemonic windows. Vertices with contributions from fewer than 10% of participants were omitted from display. Visual-area boundaries were projected to fsaverage in the same manner and displayed as group contours based on subject-overlap probability, using a 50% contour threshold. These maps were treated as descriptive visualizations of large-scale preferred-orientation organization across EVC. Using the same projected data, we also computed circular variance across participants at each fsaverage vertex to generate the complementary variability maps shown in **Supplementary Fig. 6**.

#### Retinotopic coordinates and ROI restriction

To quantify radial bias more strictly, we combined retinotopy-derived polar-angle coordinates with voxel-wise orientation tuning estimates in a V1-focused analysis (**Supplementary Figs. 4 and 5**). For each participant, voxel-wise estimates of visual-field polar angle and eccentricity were obtained from anatomy-based retinotopy outputs and matched to empirical voxel coordinates. Template-derived visual-area assignments (**Benson et al., 2014**) were compared with empirical visual-area labels as summarized in **Supplementary Fig. 3**. Because this comparison indicated the strongest correspondence in V1, inferential retinotopy-based analyses were restricted to voxels that (i) were included in the empirical EVC ROI, (ii) matched a template-derived V1 label, and (iii) had eccentricity < 8.5°. Because polar angle is defined relative to fixation, right-hemisphere angles were mirrored as 180° − ψ before pooling with left-hemisphere voxels, yielding a common 180°-periodic polar-angle axis across hemispheres. For descriptive comparison, we also summarized the distributions of mirrored polar angle and voxel preferred orientation in the sensory and mnemonic windows across this matched V1 voxel set.

#### Time-resolved orientation preferences

Voxel-wise preferred orientation over time was estimated with the same doubled-angle model described above. For each matched V1 voxel, participant, and time bin, we estimated *θ*_*pref*_(*t*) across the 12 contiguous 2-s bins spanning 4–28 s after trial onset. We additionally computed time-averaged preferences for the sensory and mnemonic windows. For the radial-bias analyses, both polar angle and preferred orientation were represented on a 180°-periodic scale so that orientations aligned with a given radial axis shared similar values.

#### Quantifying radial bias

Radial bias implies that voxels at a given polar angle prefer orientations aligned with that radial axis. To test this, we related mirrored polar angle to voxel-wise preferred orientation at each time bin. For each participant and time bin, radial bias was quantified as a circular correlation computed from mean-centered polar-angle and preferred-orientation values across matched V1 voxels. This yielded one radial-bias value per participant at each time bin. Group time courses were summarized as mean ± s.e.m. across participants, and each time bin was tested against zero with two-sided one-sample t-tests; filled symbols in the figure indicate uncorrected p < 0.05.

#### Eccentricity split

Because our stimulus (2-8.5 deg) and response dots (0.9 deg) were spatially distinguished, we repeated the same V1 radial-bias analysis separately for voxels with eccentricity < 2° and > 2° (**Supplementary Fig. 5**). Within each subset, preferred orientation was estimated with the same doubled-angle model, the same 12 time bins, and the same participant-wise circular-correlation metric.

### Choice-conditioned reconstructions

We used reconstructed orientations from the mnemonic-trained IEM to test whether trial-by-trial variability in mnemonic-period readout predicted discrimination choices and estimation bias (**Fig. 8**). For each trial, participant, and decoded test time bin, we computed a stimulus-centered decoded deviation,

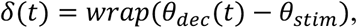

wrapped to (−90°, 90°) with 180° periodicity. Positive δ(t) indicates a CCW shift relative to the target, whereas negative δ(t) indicates a CW shift.

#### Choice-conditioned reconstructed error distributions during discrimination

We analyzed early- and late-probe discrimination trials separately using the corresponding discrimination-phase test window and restricted the analysis to the near-boundary reference offsets Δref ∈ {−4°, 0°, +4°} (**Fig. 8a**). Within each probe-timing condition, trials were further separated by discrimination choice (CCW vs CW relative to the reference). For each offset and choice condition, decoded deviations were binned in 7.5° bins, pooled with the two adjacent bins using circular wrap-around at ±90°, and renormalized to form a probability distribution. Difference panels show the pointwise difference between the CCW- and CW-conditioned distributions. The rightmost histograms summarize participant-specific decoded-deviation distributions pooled across the three near-boundary offsets separately for CCW and CW choices and then averaged across participants as mean ± s.e.m.

#### Time-resolved ROC/AUC analysis for discrimination choice

We next quantified how well decoded deviations predicted discrimination choice across the full mnemonic-IEM time course using successive 2-s test bins (**Fig. 8b**). Early- and late-probe trials were analyzed separately. Within each participant and time bin, area under the receiver operating characteristic curve(AUC/ROC) was computed separately for each near-boundary reference offset (−4°, 0°, +4°) from the ability of decoded deviations to discriminate CCW from CW choices; these AUC values were then averaged across the three offsets to obtain one participant-level AUC per time bin and probe-timing condition. AUC was computed only when both choice classes contributed at least five trials within a given offset condition. Group time courses show mean ± s.e.m. across participants. At each time bin, AUC was tested against chance (0.5) with two-sided one-sample t-tests across participants (p < 0.05, uncorrected).

#### Time-resolved ROC/AUC analysis for estimation bias

To test whether mnemonic-decoder outputs predicted continuous report variability, we analyzed early- and late-probe trials separately across the same decoded time bins (**Fig. 8c**). For each participant and each stimulus orientation, signed estimation error (reported orientation minus target orientation, wrapped to (−90°, 90°)) was median-split into CCW-biased and CW-biased classes, with the median included in both classes. Decoded deviations were then pooled across stimulus orientations after centering within 24 orientation bins (7.5° spacing, each pooling the central orientation and its two neighboring orientations), and AUC was computed at each time bin from the separation between CCW-biased and CW-biased trials. AUC was computed only when both classes contributed at least five trials. Group time courses show mean ± s.e.m. across participants, and AUC at each time bin was tested against 0.5 with two-sided one-sample t-tests across participants (p < 0.05, uncorrected).

