## Supplemental Information for "Early Visual Cortex Represents Sensory and Mnemonic Orientations in Separate Subspaces with Preserved Geometry"

**Contents**

Supplementary Note 1. Temporal isolation and prior-study timing comparison

Supplementary Note 2. dPCA validation and dimensionality robustness

Supplementary Note 3. Template matching, voxel remapping, and radial-bias analyses

Supplementary Note 4. Robustness to epoch and ROI choices

Supplementary References

These supplementary materials provide clarifying analyses and robustness checks that support interpretation of the main results, with particular emphasis on temporal isolation of late mnemonic activity, validation of the low-dimensional subspace analysis, and the relationship between mnemonic coding and coarse retinotopic radial bias.

**Supplementary Note 1. Temporal isolation and prior-study timing comparison**

The main text argues that the present design permits a more selective characterization of late mnemonic activity than is typically possible in short-delay fMRI studies of visual working memory. **Supplementary Figure 1** compares the relative timing of stimulus presentation and decoder- or encoding-model training windows across representative prior studies and the present experiment. The central issue is temporal interpretability: when model training occurs during the stimulus-evoked hemodynamic response, successful decoding from nominal delay-period activity can be difficult to attribute uniquely to actively maintained mnemonic representations rather than lingering sensory activity.

To facilitate this comparison, stimulus epochs and training windows were aligned on a common trial-time axis, and a canonical hemodynamic response function was plotted schematically to illustrate the expected temporal extent of stimulus-locked BOLD activity following a brief visual event. The hemodynamic response trace is included as an intuitive temporal reference rather than a dataset-specific fit. Under this timing framework, training windows used in many prior studies overlap more substantially with the stimulus-evoked response than does the late-delay window used here, leaving greater ambiguity about the relative contributions of sensory and mnemonic signals.

**Supplementary Figure 1** therefore clarifies the temporal rationale for the present design and analysis strategy. In the present experiment, the late-delay window was positioned to maximize separation from both stimulus-evoked activity and subsequent report-related activity, enabling a cleaner evaluation of the representational relationship between sensory and mnemonic codes in early visual cortex.

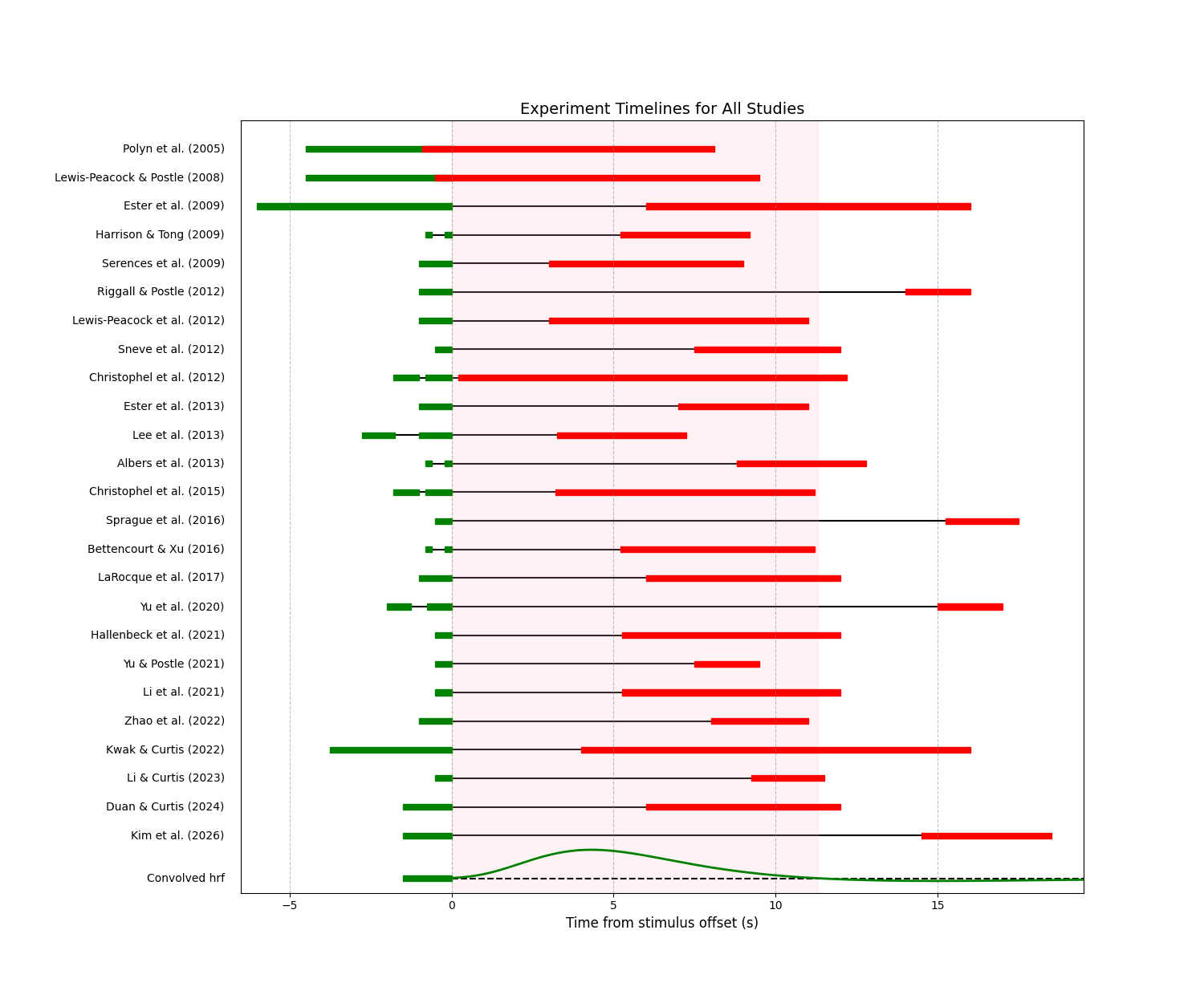
 **Supplementary Figure 1 | Timing of stimulus and decoder-training windows across prior fMRI working-memory decoding studies.**

Green bars denote stimulus presentation epochs and red bars denote decoder- or encoding-model training windows across prior fMRI studies decoding working-memory content from early visual cortex. The bottom trace shows a canonical HRF convolved with a brief stimulus (1.5 s), highlighting the temporal extent of stimulus-evoked BOLD activity (pink shading). In many prior studies, decoder training windows overlapped with the rising or falling phases of the stimulus-evoked response, complicating the separation of stimulus-driven and mnemonic contributions. In contrast, the training windows used in our work are positioned beyond the HRF tail, minimizing sensory contamination and enabling a more selective characterization of mnemonic representations.

**Supplementary Table 1.** **Timing characteristics of prior fMRI working-memory decoding studies included in Supplementary Figure 1.**

**Supplementary Table 1** summarizes the prior studies included in **Supplementary Figure 1** and makes explicit the timing assumptions used in the comparison shown there. Training windows were intended to capture the periods used to train the decoder.

| **Study** | **N** | **ROI summary** | **Stimulus epoch(s) from trial start** | **Training window from trial start** |
| --- | --- | --- | --- | --- |
| Polyn et al. (2005)^1^ | 14 | selective voxels from whole brain | 1.8-6.3 s | 5.4-14.4 s |
| Lewis-Peacock and Postle (2008)^2^ | 10 | selective voxels from whole brain (8489 voxels) | 2-6.5 s | 6-16 s |
| Ester et al. (2009)^3^ | 20 | V1 | 0-6 s | 12-22 s |
| Harrison and Tong (2009)^4^ | 6 | V1-4 | 0-0.2; 0.6-0.8 s | 6-10 s |
| Serences et al. (2009)^5^ | 7 | V1-4 | 0-1 s | 4-10 s |
| Riggall and Postle (2012)^6^ | 10 | LO and Temporal / MO | 0-1 s | 15-17 s |
| Lewis-Peacock et al. (2012)^7^ | 14 | selective voxels from whole brain, no-PFC ROI | 0-1 s | 4-12 s |
| Sneve et al. (2012)^8^ | 6 | V1-4 and LO1,2 | 0-0.5 s | 8-12.5 s |
| Christophel et al. (2012)^9^ | 19 | EVC, parietal | 0-0.8; 1-1.8 s | 2-14 s |
| Ester et al. (2013)^10^ | 20 | V1-3 | 0-1 s | 8-12 s |
| Lee et al. (2013)^11^ | 22 | lPFC, pFs, PPC | 0-1; 1.75-2.75 s | 6-10 s |
| Albers et al. (2013)^12^ | 30 | V1-3 | 0.8-1; 1.4-1.6 s | 10.4-14.4 s |
| Christophel et al. (2015)^13^ | 21 | EVC, PPC | 0-0.8; 1-1.8 s | 5-13 s |
| Sprague et al. (2016)^14^ | 6 | V1-3, IPS, sPCS | 0-0.5 s | 15.75-18 s |
| Bettencourt and Xu (2016)^15^ | 10 | V1-4, sIPS | 0-0.2; 0.6-0.8 s | 6-12 s |
| LaRocque et al. (2017)^16^ | 7 | selective voxels from whole brain | 0-1 s | 7-13 s |
| Yu et al. (2020)^17^ | 13 | V12, IPS | 0-0.75; 1.25-2 s | 17-19 s |
| Hallenbeck et al. (2021)^18^ | 7 | V1-3, LO, IPS, sPCS | 1-1.5 s | 6.75-13.5 s |
| Yu and Postle (2021)^19^ | 17 | EVC, IPS | 0-0.5 s | 8-10 s |
| Li et al. (2021)^20^ | 11 | EVC, IPS, sPCS | 0-0.5 s | 5.75-12.5 s |
| Zhao et al. (2022)^21^ | 6 | EVC | 0-1 s | 9-12 s |
| Kwak and Curtis (2022)^22^ | 11 | EVC, IPS, sPCS | 0.75-4.5 s | 8.5-20.5 s |
| Li and Curtis (2023)^23^ | 14 | EVC, IPS | 0-0.5 s | 9.75-12 s |
| Duan and Curtis (2024)^24^ | 16 | EVC, IPS | 0-1.5 s | 7.5-13.5 s |
| Kim et al. (2026) | 50 | EVC | 0-1.5 s | 16-20 s |

**Supplementary Note 2. dPCA validation and dimensionality robustness.**

The main subspace result depends on a low-dimensional representation of orientation-related activity, making it important to show that the demixed principal component analysis used in the manuscript isolates meaningful stimulus-related structure rather than an arbitrary projection. **Supplementary Figure 2** provides this validation in three ways. First, the example participant shows that condition-independent demixed components are dominated by shared task-locked temporal dynamics, whereas stimulus-related demixed components show clear separation among orientation conditions. Second, the variance summary shows that dPCA captures nearly as much total variance as conventional PCA, indicating that the demixing procedure does not discard most of the population structure. Third, the dimensionality analysis shows that the sensory-mnemonic separation is qualitatively stable across a small range of low-dimensional stimulus spaces.

In the main text, the 3D stimulus-related state space was used because it provides the minimal ambient dimensionality needed to visualize ring-like orientation structure while permitting plane-angle estimation. **Supplementary Figure 2** shows that this choice is stable: the leading stimulus-related dimensions capture the relevant orientation structure, dPCA retains nearly as much variance as PCA, and the sensory-mnemonic subspace separation remains qualitatively similar across nearby dimensionalities. These analyses support the interpretation that the plane-angle result reflects a robust property of the data rather than a narrow consequence of a single projection choice.

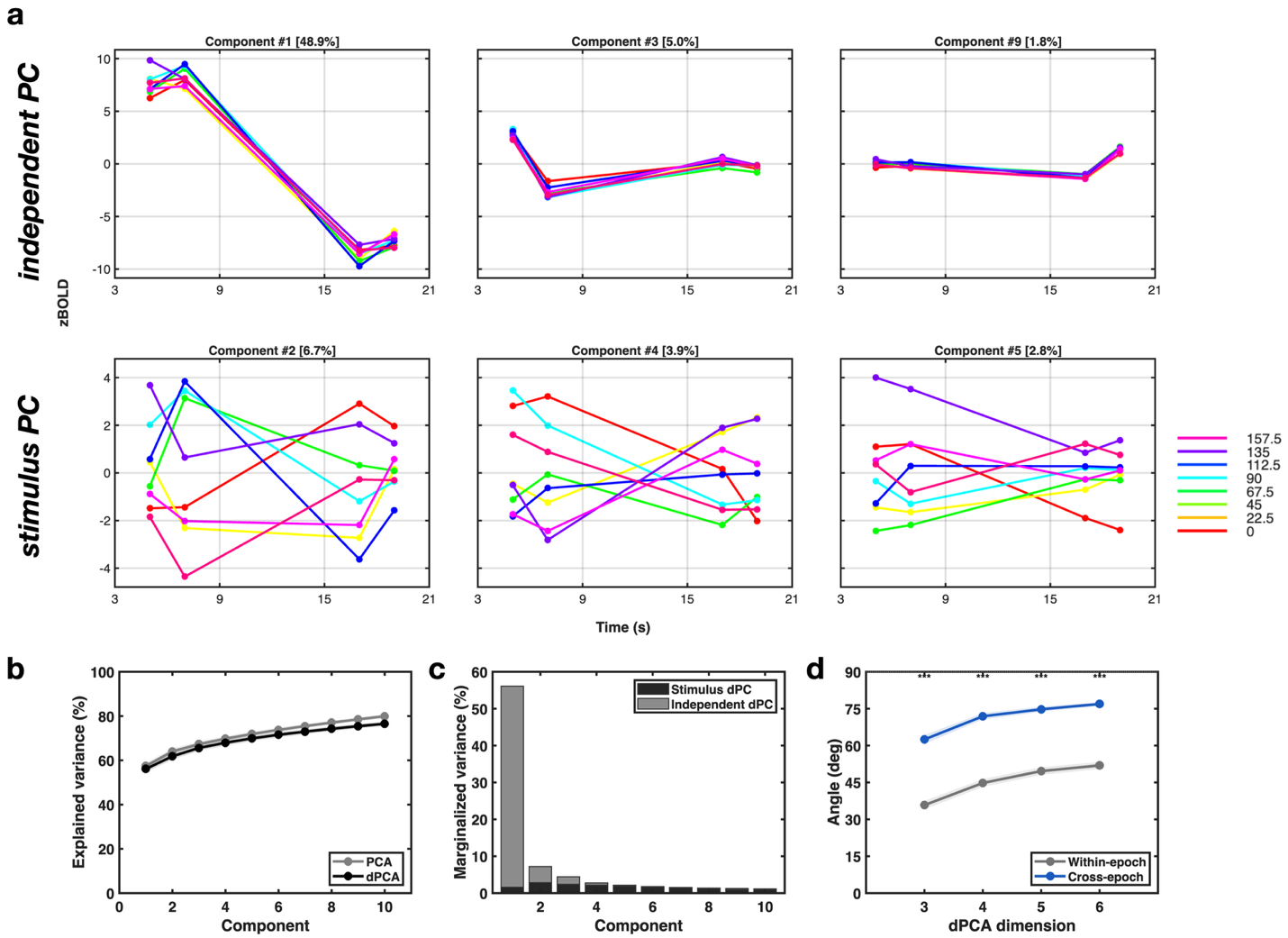
 **Supplementary Figure 2 | dPCA sanity checks and dimensionality summaries supporting the stimulus-space analysis.**

**a,** Example participant showing time courses of the first three condition-independent dPCs (top row) and the first three stimulus-related dPCs (bottom row). Colored traces correspond to the eight orientation bins. The condition-independent components were dominated by shared task-locked BOLD dynamics and showed little orientation separation, whereas the stimulus-related components showed clear orientation-dependent structure. **b,** Cumulative variance explained by the first 10 PCA components (gray) and the first 10 dPCA components (black), averaged across participants (mean ± s.e.m.). **c,** Marginalized variance explained by each of the first 10 dPCA components, stacked by marginalization. **d,** Dependence of sensory-mnemonic subspace separation on the ambient stimulus-space dimensionality. The larger principal angle between sensory and mnemonic stimulus subspaces (blue) is plotted against the within-epoch benchmark (gray). Error bands indicate mean +/- s.e.m. across participants; asterisks indicate one-sided paired tests against the within-epoch benchmark.

**Supplementary Note 3. Template matching, voxel remapping, and radial-bias analyses**

The cortical-map and voxel-preference analyses address how mnemonic orientation coding relates to voxel-wise tuning and coarse spatial organization. **Supplementary Figures 3–6** show that voxel-level preferred orientations are reorganized across sensory and mnemonic epochs, that template-based radial-bias quantification is best restricted to V1 where anatomy-based template labels show the strongest agreement with independently defined visual-area labels, and that reduced radial-bias structure during the mnemonic epoch is evident across complementary analyses.

**Note 3.1. Template matching and V1 restriction**

**Supplementary Figure 3** explains why the stricter retinotopy-based quantification was restricted to V1. Because full empirical retinotopic mapping sufficient for vertex-wise inferential analysis was not acquired for the present manuscript, anatomy-based retinotopy estimates were used to assign polar-angle and visual-area labels. The agreement analysis in **Supplementary Figure 3** shows that correspondence between independently defined visual-area labels and anatomy-based template assignments is strongest in V1 relative to V2 and V3. Template-based radial-bias quantification was therefore restricted to V1 in the inferential supplementary analyses.

**Note 3.2. Across-epoch voxel preference remapping and radial bias**

**Supplementary Figure 4** links across-epoch voxel-preference remapping to the radial-bias analysis. The reciprocal sorting analysis shows that the sensory and mnemonic epochs do not share a stable voxel-wise preference map: ordering voxels by sensory preferences highlights structure in the sensory epoch but not in the mnemonic epoch, whereas ordering by mnemonic preferences reveals the opposite pattern. This reciprocal organization is the clearest qualitative signature that the mnemonic pattern is not a simple replay, inversion, or scalar reweighting of the sensory map. The same figure then shows that, within the matched V1 voxel set used for stricter retinotopic quantification, coarse radial-bias structure is prominent during the sensory period but markedly reduced by the late mnemonic window. Thus, the mnemonic code appears both remapped at the voxel-preference level and less constrained by the coarse radial organization that characterizes sensory encoding.

**Note 3.3. Eccentricity-resolved radial-bias analysis**

**Supplementary Figure 5** addresses a natural concern about spatial specificity. In the present task, the remembered grating occupied peripheral eccentricities, whereas the response-related dot cues were near fixation. The eccentricity split therefore tests whether the reduction in mnemonic radial bias is confined to one spatial regime or remains evident across voxels covering different eccentricity ranges.

**Note 3.4. Variability of cortical-space maps**

**Supplementary Figure 6** provides a descriptive complement to the group-average cortical preferred-orientation maps shown in the main text. These variability maps contextualize the spatial consistency of the fsaverage visualizations and complement the V1-focused inferential analyses summarized in **Supplementary Figures 3–5**.

Together, **Supplementary Figures 3–6** define a consistent validation sequence: template matching motivates the V1 restriction, voxel-preference remapping shows tuning reorganization across epochs, V1-focused analyses show reduced radial-bias alignment during mnemonic coding, and cortical-space variability maps contextualize the descriptive fsaverage visualizations.

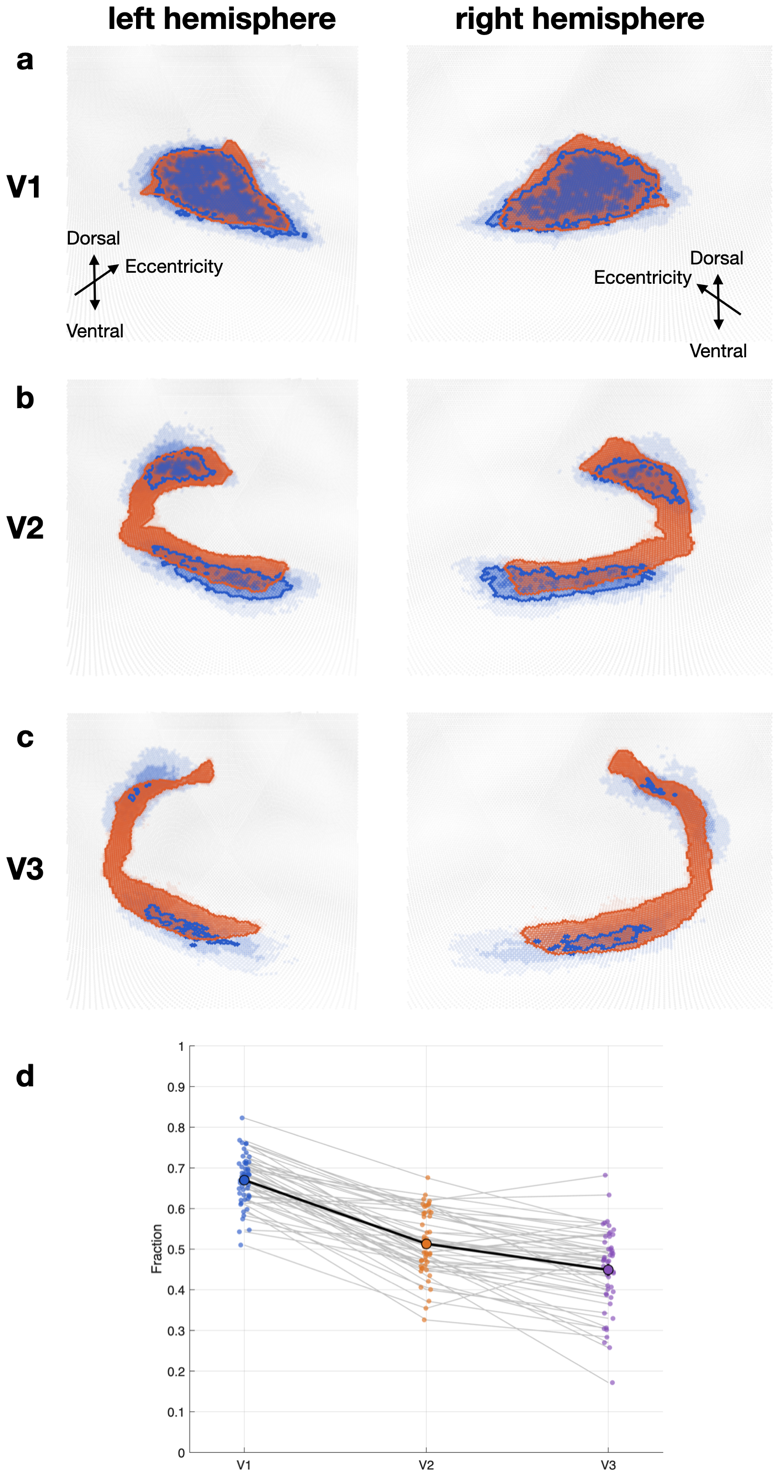
**Supplementary Figure 3 | Agreement between manually-defined and anatomy-based visual-area labels is strongest in V1.**

**a-c,** Participant-wise overlap between manually defined visual-area labels and Benson-template visual-area assignments, shown separately for V1 (a), V2 (b), and V3 (c). **d,** Overlap is summarized across participants as the fraction of manually defined voxels assigned to the corresponding anatomy-based area. Agreement was strongest in V1, providing the rationale for treating the stricter radial-bias quantification as a V1-focused analysis in the supplement rather than as an EVC-wide quantitative claim.

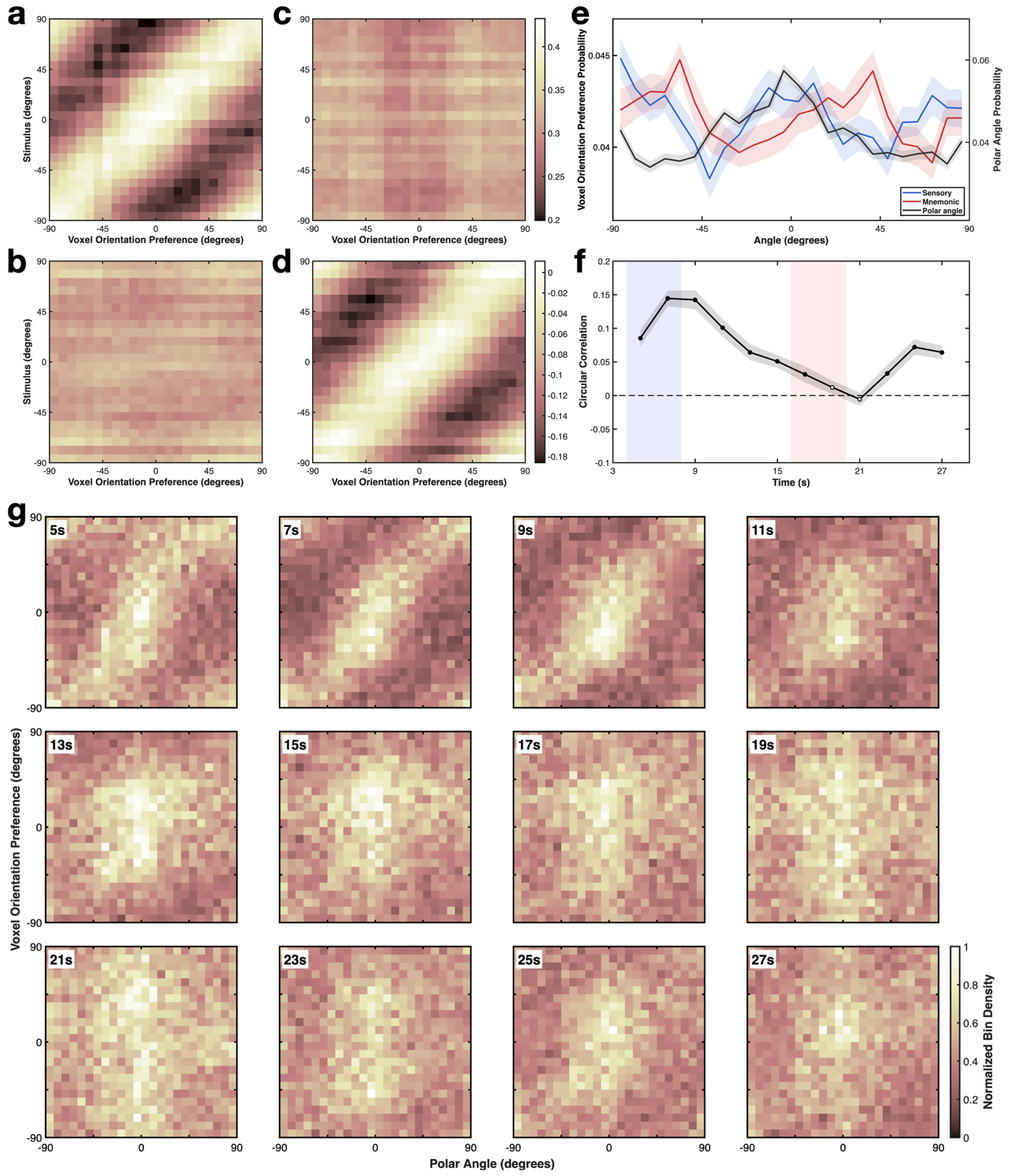
 **Supplementary Figure 4 | Voxel-level orientation preferences are remapped across epochs and V1 radial retinotopic bias is reduced during mnemonic coding.**

**a–d,** Across-epoch BOLD responses to target orientation, conditioned on voxel-wise orientation preference. Each panel plots mean BOLD response as a function of target orientation during the sensory (a,c) or mnemonic (b,d) epoch, for voxels sorted by their preferred orientation estimated in the sensory (a,b) or mnemonic (c,d) epoch. Panels a–d are based on the EVC voxel population used for the voxel-preference analysis. **e,** Distributions of voxel preferred orientations in the sensory (blue) and mnemonic (red) epochs, plotted together with the distribution of voxel retinotopic polar-angle coordinates (grey). Here, polar angle refers to the angular location of a voxel’s retinotopic position around fixation. **f,** Radial bias over time in Benson-template-defined V1 voxels, quantified as the circular–circular correlation between voxel polar angle position and preferred orientation (symbols, mean across participants; grey shading, s.e.m.; filled symbols, uncorrected p < 0.05). **g,** Time‑resolved joint distributions of voxel retinotopic polar angle (x‑axis) and preferred orientation (y‑axis) for successive time windows (4-28 s from trial onset).

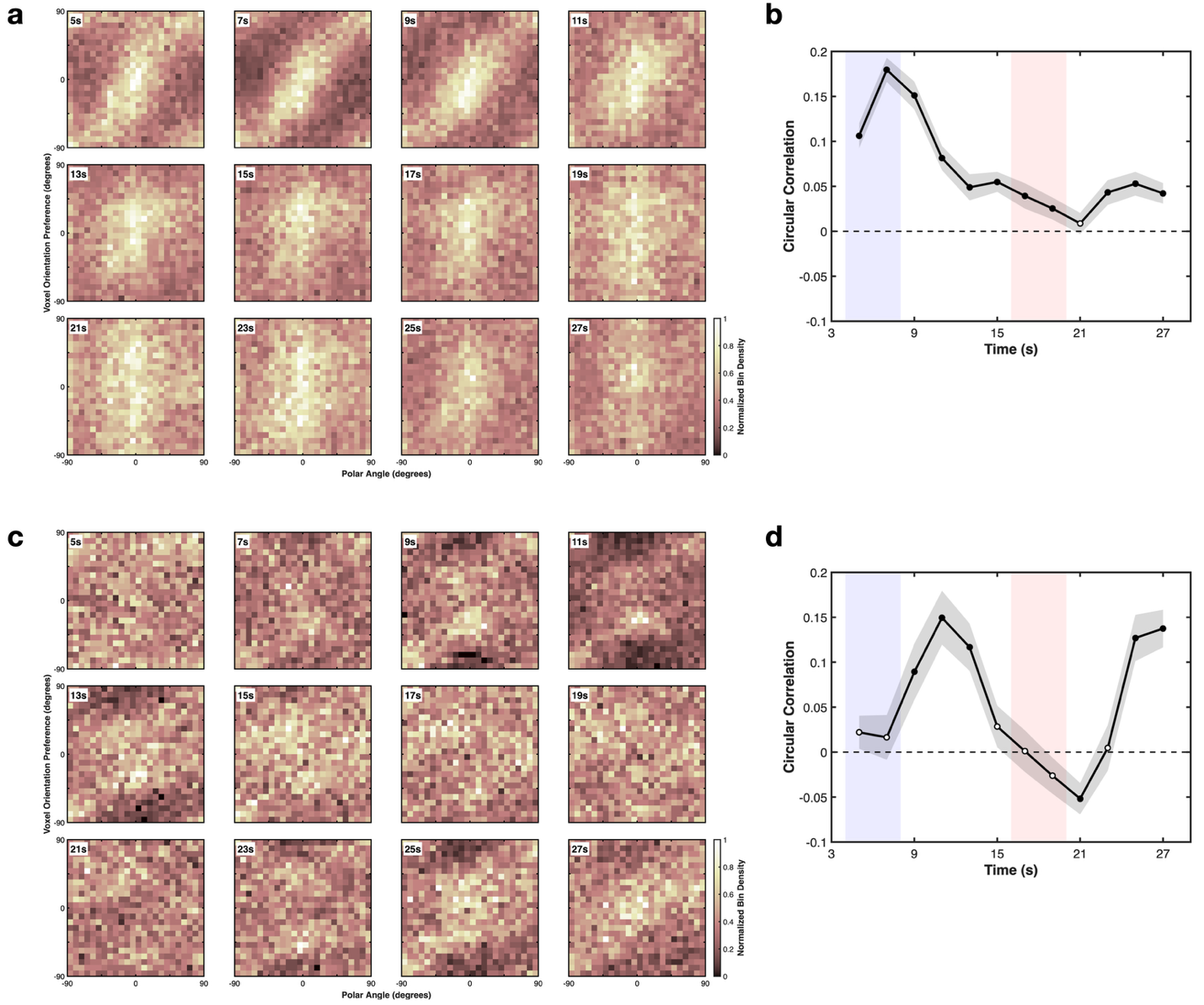
 **Supplementary Figure 5 | Time-resolved radial-bias structure in V1 shown separately for smaller and larger eccentricities.**

**a,c,** Time-resolved joint distributions of voxel retinotopic polar angle (x axis) and preferred orientation (y axis) in V1, shown separately for voxels with eccentricity < 2 degrees (a) and > 2 degrees (c). Each panel shows a 2-s time bin from 4-28 s after trial onset (labeled by bin center). Color indicates normalized bin density. **b,d,** Time-resolved circular-circular correlation between polar angle and preferred orientation for the same eccentricity groups (< 2 degrees in b; > 2 degrees in d), averaged across participants (mean +/- s.e.m.). Blue and red shaded regions denote the sensory (4-8 s) and mnemonic (16-20 s) epochs, respectively; filled markers indicate uncorrected p < 0.05 against zero. Radial organization was weak and variable at smaller eccentricities, but was clearer at larger eccentricities early in the trial. In both eccentricity ranges, the correlation was low around the late mnemonic window, consistent with reduced radial-bias structure during mnemonic coding relative to the sensory period.

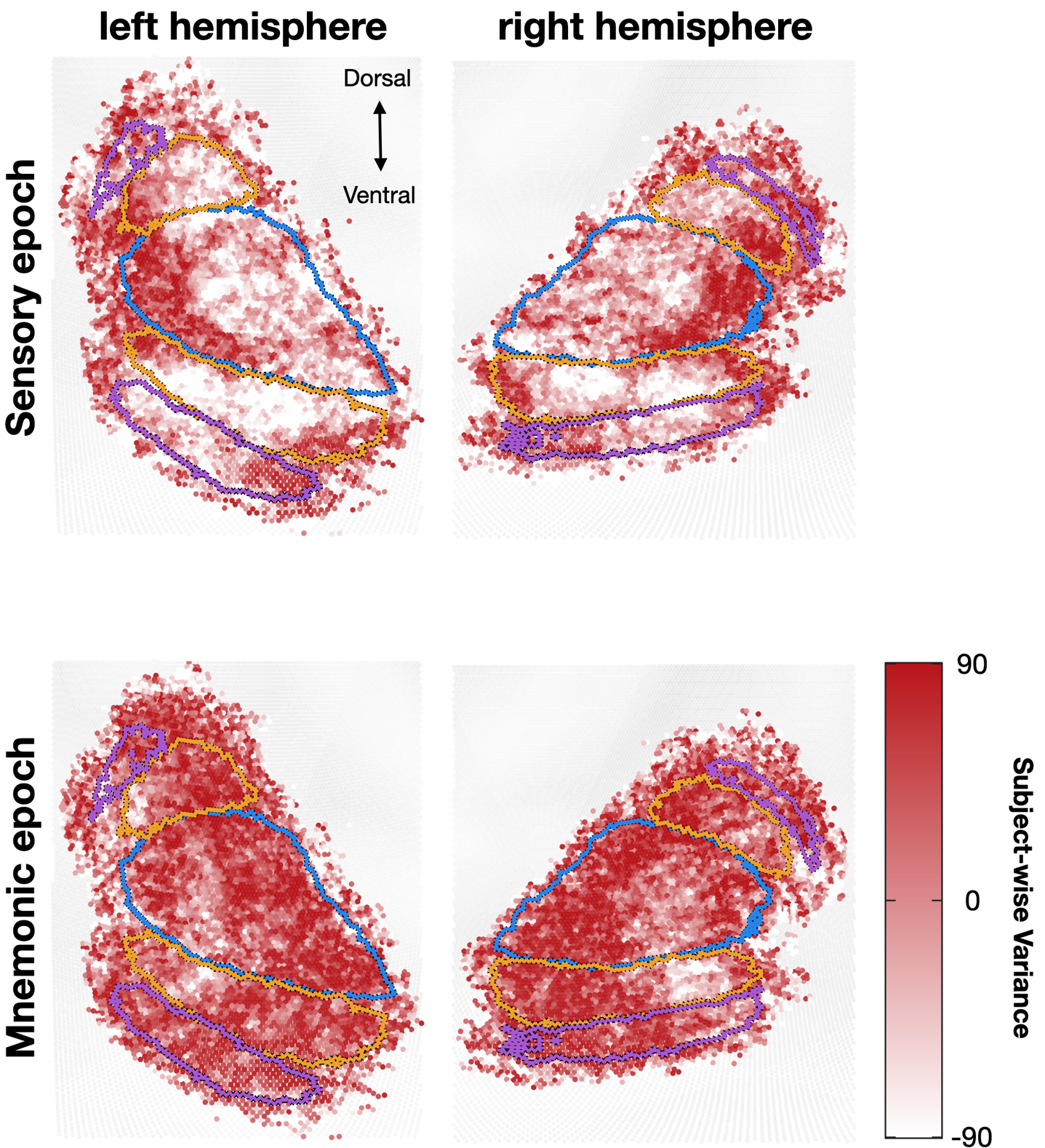
 **Supplementary Figure 6 | Across-participant variability of preferred-orientation maps in cortical space across sensory and mnemonic epochs.**

Cortical-space maps showing across-participant variability of preferred-orientation estimates on fsaverage flatmaps for the left and right hemispheres. **Top,** sensory epoch. **Bottom,** mnemonic epoch. Overlaid contours delineate visual-area boundaries across early visual cortex. These maps provide a descriptive visualization of the spatial distribution of variability in preferred-orientation estimates across epochs.

**Supplementary Note 4. Robustness to epoch and ROI choices**

**Supplementary Figures 7 and 8** test whether the main conclusions are robust to two reasonable analysis choices: the precise definition of the mnemonic window and the pooling of V1, V2, and V3 into a single EVC ROI.

**Note 4.1. Robustness to a stricter mnemonic window**

**Supplementary Figure 7** shows that the main IEM, dPCA, and RSA patterns are preserved when the mnemonic epoch is defined more conservatively, excluding the 18–20 s interval nearest the estimation period. The qualitative pattern remained similar when the sensory and mnemonic analyses were repeated using 4–6 s and 16–18 s windows. These analyses indicate that the main sensory-mnemonic differences do not depend on inclusion of the 18–20 s interval.

**Note 4.2. Robustness to ROI pooling**

**Supplementary Figure 8** shows that the core findings are reproduced when V1, V2, and V3 are analyzed separately rather than pooled into a single EVC ROI. Because voxel counts decreased systematically from V1 to V3, apparent magnitude differences across subareas were not interpreted as formal evidence for hierarchical differences.

Together, **Supplementary Figures 7 and 8** show that poor cross-epoch generalization, separable low-dimensional subspaces, and preserved within-epoch orientation geometry are robust to stricter timing and ROI definitions.

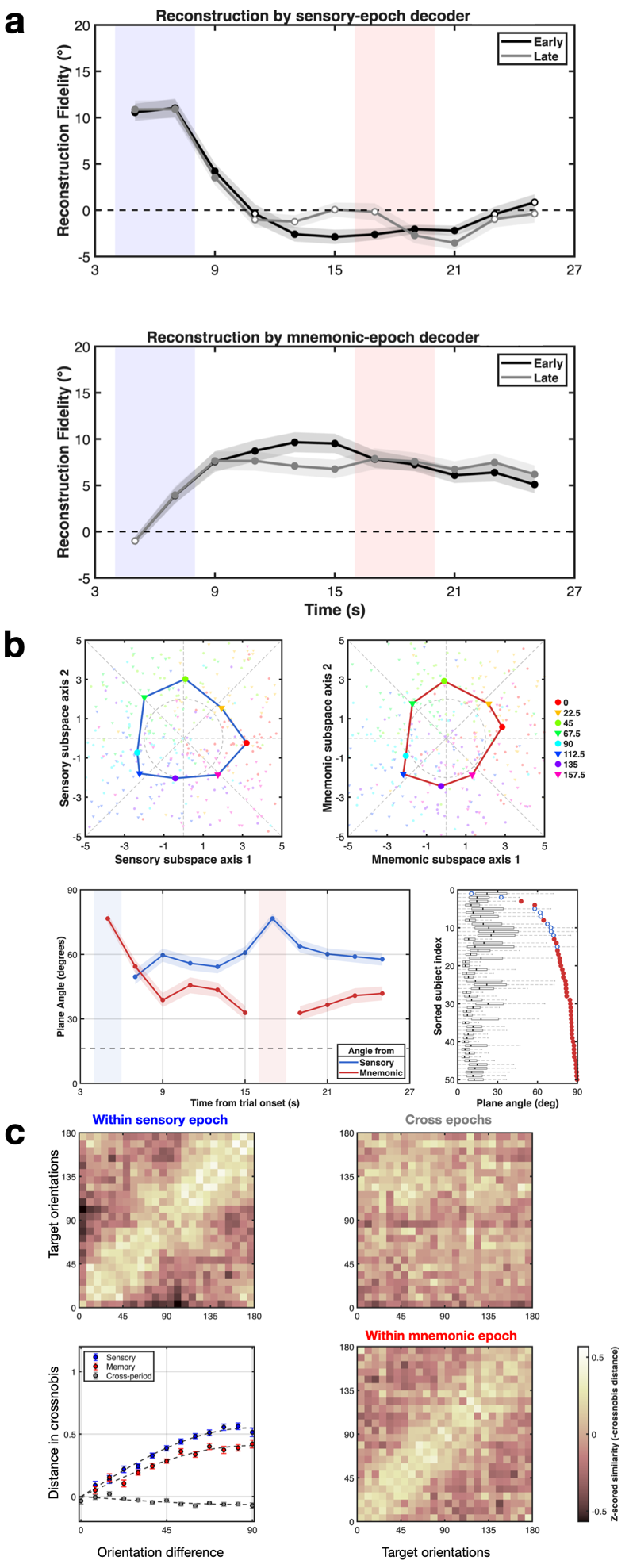
**Supplementary Figure 7 | Core IEM, dPCA, and RSA results are reproduced using a stricter mnemonic epoch definition.**

**To address the possibility that the mnemonic epoch used in the main analyses (16–20 s) could be influenced by the estimation period beginning at 18 s, we repeated the key analyses using a sensory window at 4–6 s and a mnemonic window at 16–18 s. a, IEM reconstruction fidelity reproduced the main pattern shown in Fig. 3: decoding trained on the sensory epoch peaked early and declined across the delay, whereas decoding trained on the mnemonic epoch remained robust throughout the delay. b, dPCA reproduced the main geometric structure shown in Fig. 4, including ring-like sensory and mnemonic manifolds and a clear separation between sensory and mnemonic subspaces across time and participants. c, RSA reproduced the main representational geometry shown in Fig. 5: orientation-dependent structure was evident within the sensory and mnemonic epochs, whereas cross-epoch similarity remained comparatively weak. Together, these results show that the main findings do not depend on including the 18–20 s interval in the mnemonic epoch.**

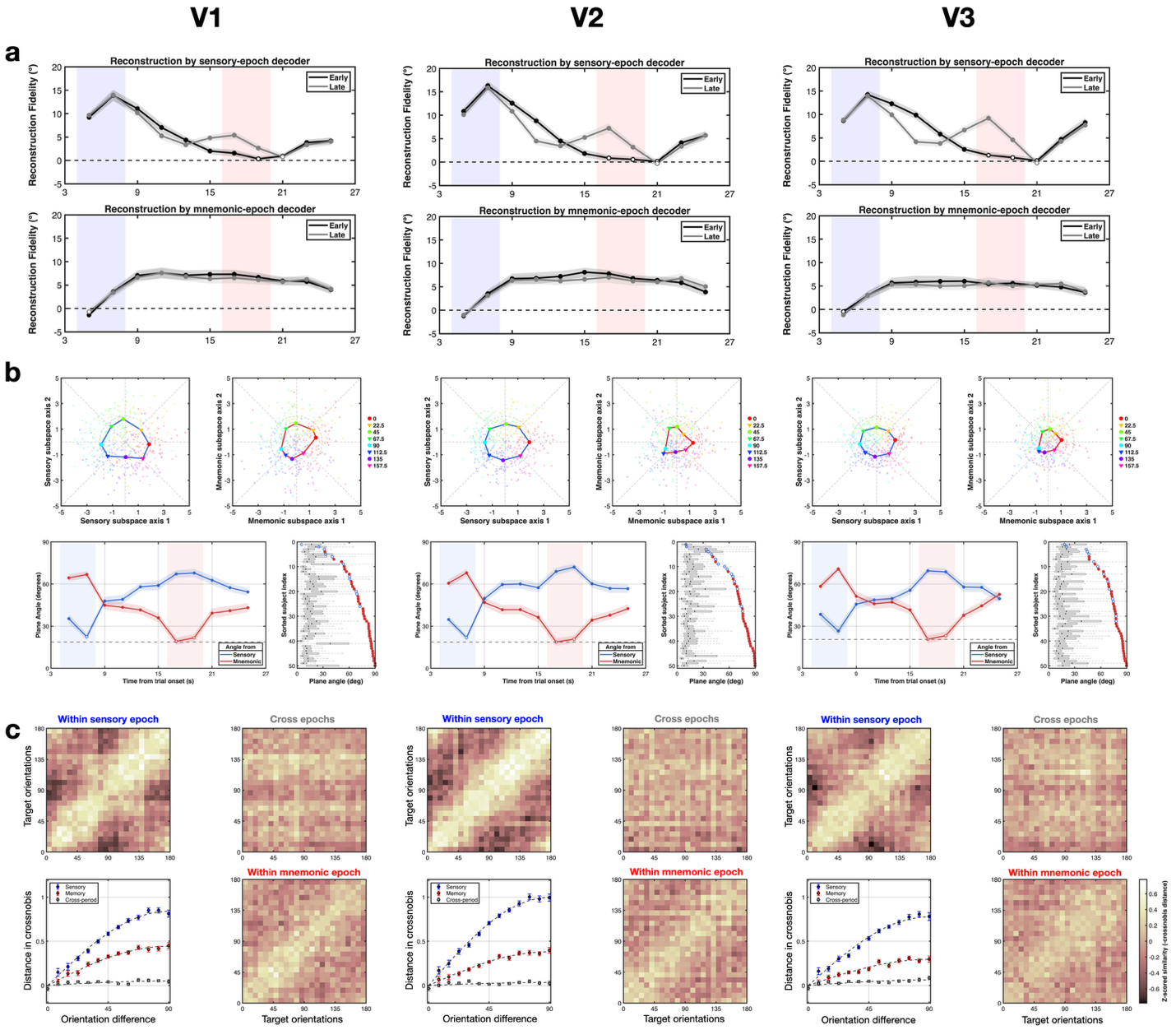
 **Supplementary Figure 8 | Core IEM, dPCA, and RSA results are reproduced when V1, V2, and V3 are analyzed separately.**

To address the concern that the main results might depend on pooling V1, V2, and V3, we repeated the key analyses separately for each visual subarea. Columns show results for V1, V2, and V3, respectively. **a,** IEM results reproduced the main pattern shown in **Fig. 3** in each subarea: reconstruction trained on the sensory epoch was strongest early in the trial, whereas reconstruction trained on the mnemonic epoch remained robust during the delay period. **b,** dPCA results reproduced the main geometric structure shown in **Fig. 4** in each subarea, including ring-like sensory and mnemonic manifolds, temporal separation between sensory and mnemonic subspaces, and consistent plane-angle structure across subjects. **c,** RSA results reproduced the main representational geometry shown in **Fig. 5** in each subarea, with clear orientation-dependent similarity structure within the sensory and mnemonic epochs and comparatively weaker cross-period similarity. Although reconstruction fidelity, manifold size, and plane-angle separation may appear to decrease from V1 to V3, these trends should not be interpreted as quantitative differences between subareas, because the number of voxels also decreased systematically across areas. Mean voxel counts (mean ± s.e.m., n = 50 subjects) were 597.4 ± 19.5 for V1, 438.6 ± 15.4 for V2, and 310.6 ± 11.2 for V3. Together, these analyses show that the central findings are not driven by combining V1, V2, and V3 into a single pooled visual-cortical region.
